# Improved chromatin quantitative trait loci mapping with CACTI

**DOI:** 10.1101/2025.06.06.657866

**Authors:** Lili Wang, Zining Qi, Xuanyao Liu

## Abstract

Mapping chromatin quantitative trait loci (cQTLs) is crucial for elucidating the regulatory mechanisms governing gene expression and complex traits. However, current cQTL mapping methods suffer from limited detection power, particularly at existing sample sizes, and are constrained by peak-calling accuracy. To address these limitations, we present CACTI, a novel method that improves cQTL mapping by leveraging correlations between neighboring regulatory elements. Across diverse histone marks (H3K4me1, H3K4me3, H3K27ac, H3K27me3 and H3K36me3) and cell types, CACTI identifies 51%-255% more cQTL signals compared to traditional single-peak-based approaches. Using CACTI, we generate a comprehensive cQTL map for the five histone marks across multiple cell types and perform colocalization analyses with GWAS loci from 44 complex traits. CACTI cQTLs colocalize with 6%-47% of GWAS loci, which is on average 15%-424% more than the standard cQTL mapping method across different marks. 24%-75% of colocalized GWAS loci show no colocalization with eQTLs. This underscores CACTI’s unique ability to uncover regulatory mechanisms that would otherwise remain undetected by eQTL analysis alone, significantly improving the functional interpretation of GWAS findings.

## INTRODUCTION

The majority of disease-associated loci identified by genome-wide association studies (GWAS) are found in non-coding regions of the genome^1^. They have been found to be enriched for genetic variants influencing gene regulation (i.e., expression quantitative trait loci or eQTLs), suggesting their regulatory functions^2^. Yet, variations in steady-state gene expression only account for a small proportion of trait heritability^3–6^. The missing link has led to a growing interest in exploring additional molecular traits that may help explain the regulatory mechanisms underlying GWAS loci. One particularly promising area of study is the regulatory impact of **chromatin structure**, the local molecular organization of the genome resulting from the association of DNA fibers with histone proteins. Changes in chromatin organization can alter the accessibility of genomic sequence elements to regulatory molecules such as transcription factors (TFs). As such, they can exert context-dependent effects on gene expression that may not be detectable in non-relevant cell types or by traditional eQTL mapping methods that rely on measures of steady-state transcriptional activity.

Recent advancements in high-throughput assays have opened up new opportunities for studying chromatin structure and its mechanistic impact on complex traits^7^. Many studies employing these emerging methods have identified genomic locations where allelic variation influences chromatin state, as characterized by histone modifications (histone QTLs), TF binding (footprint QTLs), or chromatin accessibility (chromatin accessibility QTLs) throughout the genome^8–12^. We collectively refer to these loci as chromatin QTLs (cQTLs). Other studies have demonstrated that cQTLs can exert “priming effects”^13–15^, where steady-state genetic effects on the chromatin structure of nearby regulatory elements can drive context-dependent changes in gene expression. For example, Baca et al. showed that cQTLs can reveal variants associated with prostate cancer risk that are not observed as eQTLs, likely because their influence on disease is mediated through transcription factor signaling under specific biological conditions that are not represented in steady-state expression datasets^15^. Such findings indicate that mapping cQTLs can provide critical insights into how genetic variation impacts chromatin state, gene regulation, and complex traits.

Nevertheless, two persistent challenges compromise cQTL detection. First, most epigenetic studies have limited sample sizes^9,14^, reducing their statistical power to detect cQTLs. Second, accurate peak calling is difficult for certain histone modifications^16^, particularly those covering broad or mixed regions with lowly enriched read coverage^17^. These modifications often yield inconsistent peak calls depending on the performance of peak callers^16^. These challenges hinder our ability to capture a complete map of cQTLs and underscore the need for more powerful and robust methods to improve detection power and accuracy in cQTL mapping.

To address these challenges, we introduce CACTI, a method that identifies cQTLs by leveraging correlations in measures of chromatin conformation among nearby *cis*-regulatory elements. Studies show that regulatory elements in close proximity often exhibit coordinated regulation and share genetic control, likely influenced by the three-dimensional organization of the genome^18,19^. CACTI incorporates these correlations into the cQTL mapping process using a multivariate association test^20^, which detects cQTLs of coregulated elements jointly. Multivariate association methods tend to be more powerful than traditional univariate methods, enhancing cQTL detection power and uncovering broader genetic influences on chromatin regulation. Additionally, we introduce segment-based CACTI (CACTI-S), which skips peak calling entirely and maps cQTLs directly from sequencing reads, eliminating the bias of peak calling introduced for certain peak structures.

We applied CACTI to a collective dataset of various epigenetic histone marks known to alter chromatin structure (H3K27ac, H3K4me1, H3K4me3, H3K36me3, and H3K27me3) in several cell types and tissues (LCL, macrophage, brain, heart, lung, and muscle) and constructed a comprehensive map of associated cQTLs. CACTI consistently outperformed traditional single-peak–based mapping methods for marks with either narrow or broad peak signatures. Furthermore, our map of cQTLs provided insights into the regulatory mechanisms underlying trait-associated loci.

## RESULTS

### Overview of the method

We developed a method, called CACTI, to increase the power of chromatin QTL (cQTL) detection. In contrast to standard cQTL calling methods that test the associations between single nucleotide polymorphisms (SNPs) and individual peaks from chromatin structure assays, CACTI leverages the correlation structure among proximal peaks^18,19^ by using multivariate association tests^20^ to explore the correlations between a given SNP and multiple nearby peaks. This approach provides substantially increased power to detect cQTLs. The method consists of two main steps (Figure 1).

**Figure 1.**
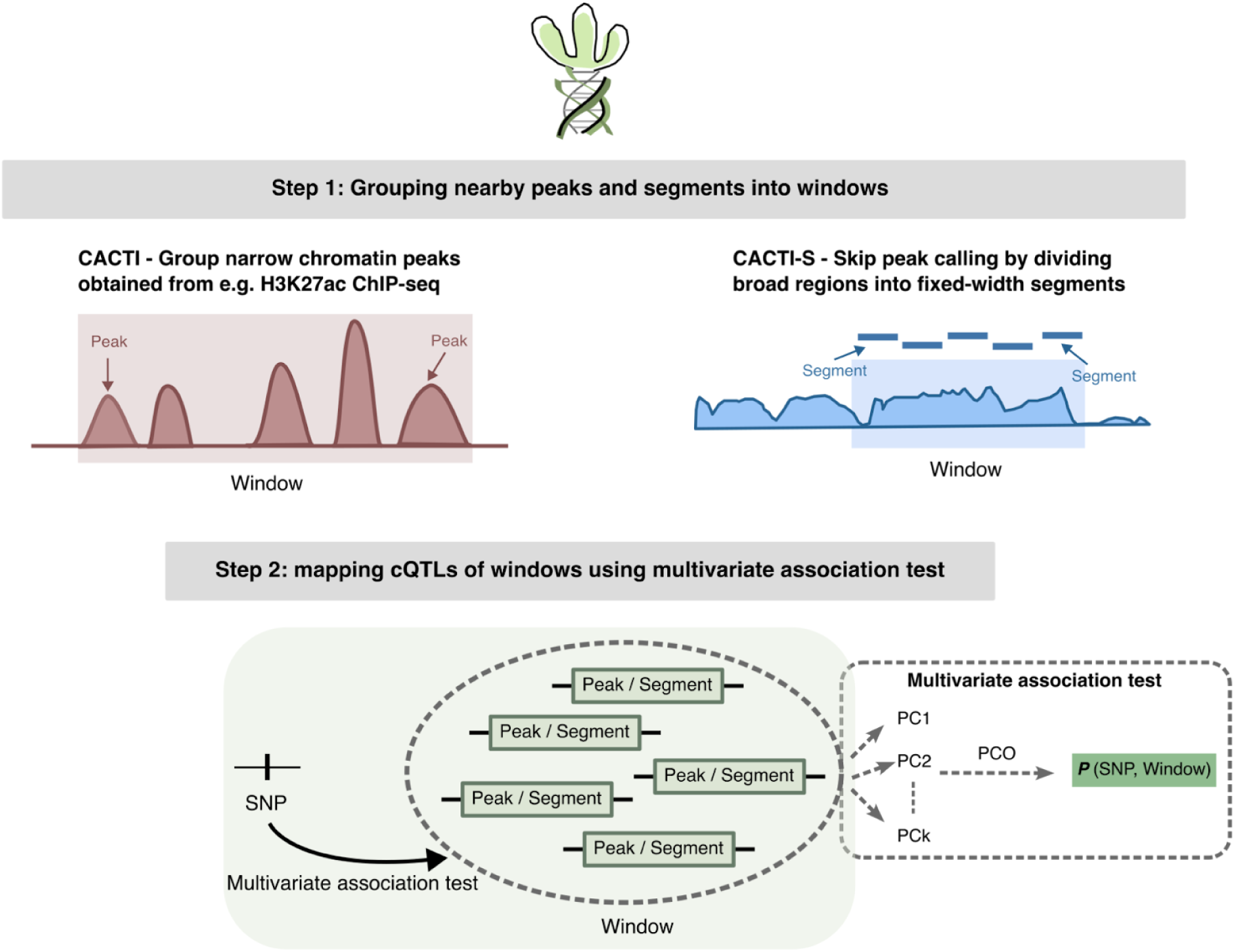
Overview of CACTI. CACTI identifies cQTLs by jointly modeling physically adjacent and correlated regulatory signals within fixed-width genomic windows. The method consists of two main steps. **Step 1:** nearby correlated chromatin modification or accessibility peaks are grouped into non-overlapping windows using a fixed-width window strategy. For histone marks where peak calling is unreliable, we apply a segment-based strategy, CACTI-S, which partitions regulatory regions into fixed-width segments and groups these segments into windows, thereby avoiding reliance on peak calling. **Step 2:** within each window, correlated peaks/segments are decomposed into principal components (PCs), and genetic association is tested using the PC-based omnibus (PCO) test to combine signals across components.

In Step 1, CACTI groups neighboring peaks from assays measuring histone modifications (e.g., ChIP-seq) or chromatin accessibility (e.g., ATAC-seq) within fixed-width genomic windows. For histone marks with narrow peak signatures, such as H3K27ac, H3K4me3, and H3K4me1, the default window size is 50 kb. This window size is sufficiently large to include multiple peaks in most windows, enabling effective aggregation of correlated signals, while remaining small enough to avoid diluting shared regulatory effects. CACTI groups peaks within consecutive genomic intervals (e.g., 0–50 kb, 50–100 kb), using the start position of the first and last peaks to define the window boundaries. Peaks spanning multiple windows are assigned to the first window they overlap. We performed several secondary analyses to assess the robustness of CACTI to window definitions. First, we evaluated performance across multiple window sizes (10 kb, 25 kb, and 50 kb) for H3K27ac, H3K4me3, and H3K4me1, and observed that cQTL detection power is largely insensitive to window size (Supplementary Notes; Figure S1). Second, we shifted all window start positions by 25 kb and found that the majority of cQTL signals were replicated before and after shifting (Supplementary Notes; Figure S2), indicating that CACTI results are stable and not sensitive to the initial window placement.

In Step 2, CACTI tests associations between multiple peaks in each window and *cis* SNPs (i.e., those within the window and the regions closely flanking it). The default *cis* region size is 100 kb, but can be adjusted depending on the parameters of the study. Association testing is performed using a powerful and robust Principal Component (PC)-based omnibus test (PCO)^20,21^. PCO is a multivariate test that transforms correlated peaks into orthogonal PCs and combines six PC-based statistical tests—PCMinP, PCFisher, PCLC, WI, Wald, and VC—to achieve robustness under unknown genetic architectures while maintaining a high statistical power (Methods). Since multiple PCs are likely to contain association evidence, and it is difficult to predict which PCs have the largest power to identify cQTLs, PCO utilizes all PCs. Each PC-based test combines multiple PCs differently; for example, PCLC linearly combines each PC weighted by eigenvalues, and Wald quadratically combines the PCs (Methods). Using all six PC-based tests enables capture of signals under a variety of genetic architectures. For each window, the PCO test statistic is defined as

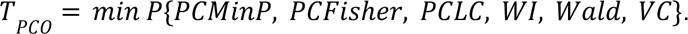

The association p-value can be computed by performing an inverse-normal transformation of the test statistic and is efficiently computed in CACTI using a multivariate normal distribution (Methods). For windows containing only one peak, p-values are obtained using the standard univariate association test. To adjust for multiple testing, we estimate the false discovery rate (FDR) for each window tested—regardless of the number of peaks within—by calculating Storey’s q-value^22^ for the SNP most strongly associated with that window. We refer to significant windows with at least one associated cQTL SNP as cWindows.

For histone marks that have broad peak signatures and low signal-to-noise ratios (such as H3K36me3 or H3K27me3), it can be difficult to accurately call peaks and map cQTLs^16^. We therefore developed segment-based CACTI (CACTI-S), which maps cQTLs directly from the read counts of sequencing-based chromatin structure assays without peak calling. Specifically, instead of calling peaks, CACTI-S divides the genome into small non-overlapping segments (e.g., 5 kb), filters out the low-quality segments with extremely low counts of aligned reads, and groups the remaining segments using non-overlapping fixed-width windows (Methods). Similar to CACTI, this method uses PCO to test the association between SNPs and the window regions containing multiple segments. CACTI-S leverages correlations between neighboring segments and further increases detection power by mitigating the impact of inaccurate peak calling on cQTL mapping.

### CACTI substantially outperforms single-peak–based mapping

To evaluate the performance of CACTI, we analyzed ChIP-seq data of the histone mark H3K27ac from lymphoblastoid cell lines (LCLs) of 78 Yoruba individuals^9,23^. We grouped a total of 100,844 peaks called in Fair et al.^23^ into non-overlapping 50 kb windows, each of which contains 1-21 peaks. In total, CACTI identifies 7027 cWindows (25% of total tested windows), which are windows with at least one significant cQTL SNP within the defined *cis* region at 5% FDR. For this test, we set a 500 kb *cis* region to be consistent with the original study^23^.

We compared CACTI to the standard single-peak–based method implemented by QTLtools^23,24^ by examining the overlap between cWindows detected by CACTI and cPeaks with at least one cQTL SNP identified by single-peak–based mapping. To allow a meaningful comparison of the number of signals between CACTI and the single-peak method, we converted the number of cPeaks to an equivalent number of windows, by counting the number of windows containing at least one cPeak. CACTI identifies significantly more cQTL signals than the single-peak method: QTLtools detects 4967 cPeaks, 96% of which map to 3577 cWindows detected by CACTI, while CACTI detects an additional 3450 cWindows (49% of total cWindows; Figure 2A). Notably, this increase in discovery occurs even though the *cis* regions tested by CACTI are more stringently defined and typically smaller compared to those used by the single-peak univariate test, because CACTI requires tested variants to fall within the defined *cis* region of all peaks within a window.

**Figure 2.**
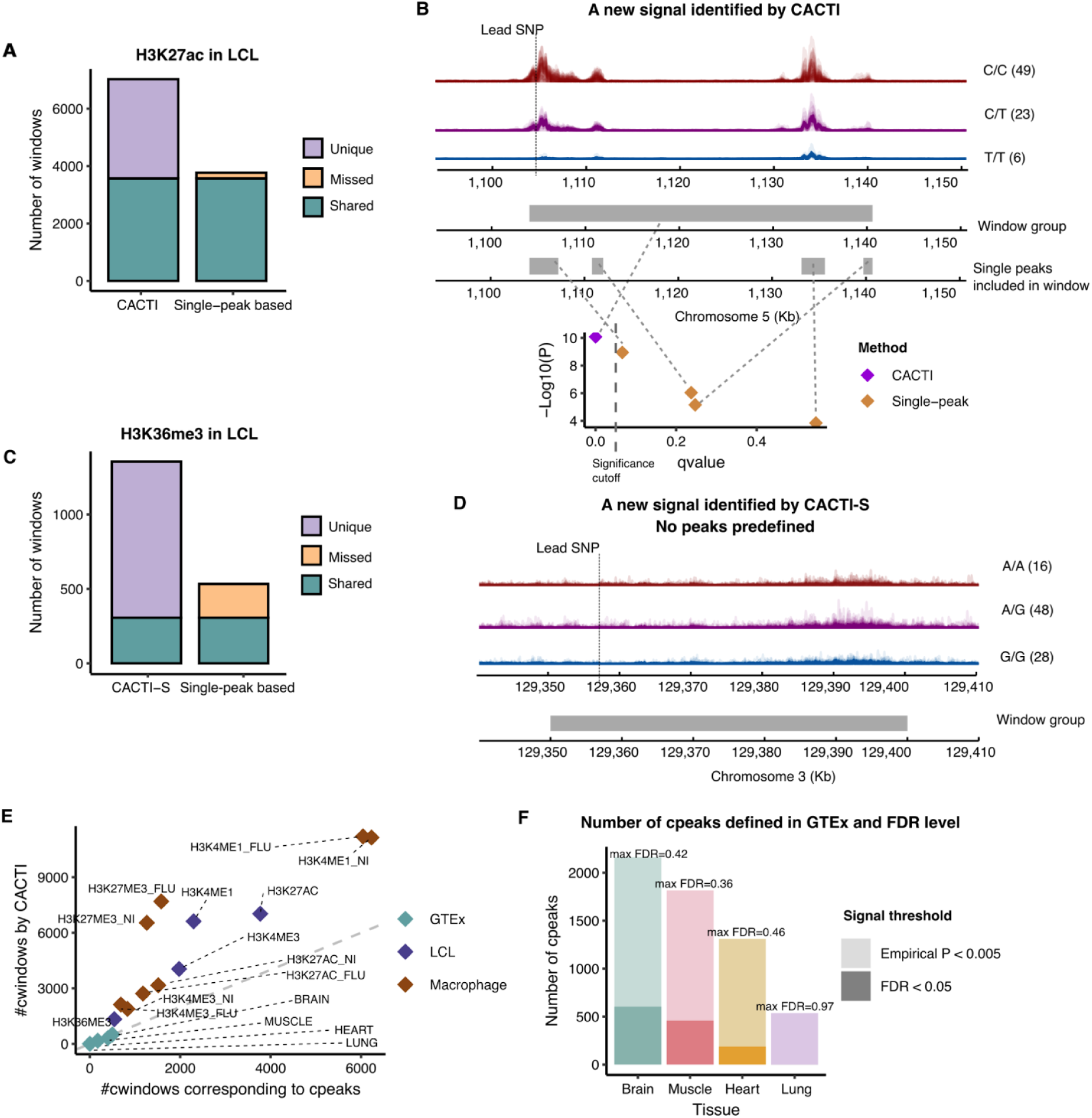
Performance comparison of CACTI versus standard single-peak–based methods. (A) Comparison of signals detected by CACTI versus single-peak–based mapping method for the mark H3K27ac in LCL. Shared cQTL signals are represented in green, signals specific to single-peak–based methods (i.e., missed by CACTI) are represented in yellow, and signals specific to CACTI (i.e., missed by single-peak methods) are represented in purple. The number of cPeaks was converted to the number of cWindows to facilitate fair comparisons. (B) An example of a new cWindow signal discovered by CACTI that is undetected by the single-peak method. The upper track displays the aligned H3K27ac reads for individuals stratified by the genotypes of the lead cQTL in this region (marked by a dashed line). The middle track represents the cWindow region. The bottom track shows the four single peaks included in the window. The scatter plot shows the p-value and q-value from association tests of the window and single peaks. (C) Comparison of signals detected by CACTI-S versus single-peak–based mapping method for the mark H3K36me3 in LCL. Shared cQTL signals are represented in green, signals specific to single-peak–based methods (i.e., missed by CACTI) are represented in yellow, and signals specific to CACTI (i.e., missed by single-peak methods) are represented in purple. The number of cPeaks was converted to the number of cWindow to facilitate fair comparisons. (D) An example of a CACTI-S cWindow, where no peaks were confidently called. The upper track displays the aligned H3K36me3 reads for individuals stratified by the genotypes of the lead cQTL in this region (marked by a dashed line). The middle track represents the cWindow region. (E) Comparison of the number of signals identified by CACTI and standard mapping methods across various datasets. Y-axis shows the number of cWindows identified by CACTI. X-axis shows the number of windows containing at least one cPeak identified by the standard single-peak–based mapping. Each dot represents a histone mark. Color represents different datasets. (F) The number of cPeak signals in eGTEx under the p-value cutoff used in the original paper (empirical p-value<0.005) and under a more stringent p-value cutoff (FDR<0.05). The X-axis is the four eGTEx tissues. The Y-axis is the number of cPeaks defined under two different p-value thresholds. The darker color represents cPeaks defined with multiple testing correction, under FDR<0.05. The lighter color represents cPeaks using the p-value threshold used in the eGTEx paper^10^, i.e., empirical p-value < 0.005. Labels on the top of the bars show the maximum FDR levels of cPeaks defined under the empirical p-value threshold.

To validate cQTLs uniquely detected by CACTI, we examined their enrichment for blood eQTLs from a large eQTLGen Consortium dataset^25^. We expect that bona fide cQTLs—particularly those regulating H3K27ac—should also exert measurable effects on gene expression due to their impact on chromatin accessibility. While many of these effects will be context-driven and thus difficult to discover by typical eQTL mapping methods, a higher-than-chance proportion of cQTLs should still be detected in large-scale eQTL datasets like this one. Indeed, we found that CACTI-specific cQTLs were significantly enriched in eQTLs (enrichment odds ratio (OR)=1.39, p-value = 1.07x10^-12^; Figure S3, Table S2, Methods), suggesting that the novel cQTLs likely have regulatory relevance. As a secondary analysis, we re-evaluated eQTL enrichment using background windows matched on the number of aggregated peaks to control for potential biases arising from differences in peak density. Enrichment was assessed using a mixed-effects logistic regression model with a random intercept for each matched set. Under this peak-density–matched design, CACTI-specific cQTLs remained significantly enriched for eQTLs (enrichment OR = 1.36, p-value=1.63x10^-11^; Table S3, Methods), indicating that the observed enrichment is not driven by differences in peak density among cWindows.

CACTI outperforms single-peak mapping primarily due to its ability to capture signals with poorer signal-to-noise ratios, which can be quantified by z-scores of cQTLs. Specifically, the univariate association z-scores of cQTLs identified by CACTI are significantly smaller than those of cQTLs by the single-peak mapping method (comparison p-value = 2.2x10^-16^; two-sided t-test; Figure S4), indicating that CACTI is more sensitive to weaker genetic effects with lower signal-to-noise ratios. This improvement confirms our expectation that CACTI’s ability to leverage correlated signals among neighboring peaks should allow it to detect subtle but consistent genetic effects that are missed by univariate testing. For example, CACTI identifies cQTLs associated with a window on chromosome 5 containing four peaks (lead p-value = 8.35x10^-11^), while single-peak mapping detects no significant associations with this region (p-values = 1.13x10^-9^ to 1.43x10^-4^ for each single peak; Figure 2B).

To further verify that the additional signals identified by CACTI arise from increased statistical power rather than inflated false positives, we performed permutation analyses to assess T1E rates. We permuted sample labels and peak labels. Across 10 independent permutations, the T1E rate is 0.012 at 1% and 0.049 at 5% (Methods, Figure S5), supporting that CACTI is well controlled for false positives. As a secondary analysis, we assessed the replication rates of CACTI signals by randomly splitting the samples into two subsets of equal sample sizes (N=36, Supplementary Notes). We applied both CACTI and single-peak–based methods to both subsets, and used a replication p-value threshold of 0.05 divided by the total number of signals. Although the reduced sample size (N<50) substantially decreased the number of detectable signals (which is expected but limits overall power and constrains replication rates), CACTI consistently exhibited higher replication than single-peak–based methods, with 1.45- to 2.96-fold increases (Figure S5). Replication rates are expected to improve with larger sample sizes, although such data are not currently available for evaluation.

### CACTI-S outperforms single-peak–based mapping for broad mark H3K36me3 without peak calling

Some histone marks, such as H3K36me3, exhibit broad enrichment patterns with low signal-to-noise ratios, making peak calling challenging and reducing cQTL detection power. To overcome this limitation, CACTI-S bypasses peak calling and directly maps cQTLs by testing high-quality read segments grouped into fixed-size windows (Methods).

To evaluate the performance of CACTI-S, we analyzed a CUT&TAG dataset of the histone mark H3K36me3 in LCLs^23^. We divided the genome into small 5 kb segments, filtering out any segments with no or extremely low read alignments to reduce noise (Methods), and grouped the remaining segments into 50 kb fixed-width windows. We tested the associations between the window regions and *cis* variants (within 500 kb) using the multivariate approach^20^ described above (Methods). In total, CACTI-S detects 1355 cWindows (4.82% of total tested windows) with at least one cQTL SNP at 5% FDR (Figure 2C). We compared the performance of CACTI-S with the standard single-peak-based cQTL mapping method described in Fair et al.^23^ (Methods). The single-peak mapping method identifies only 338 cPeaks (2.41% of the total peaks tested) associated with at least one cQTL SNP (Figure 2C). Most of the cPeaks (246/338; 73%) overlap with cWindows detected by CACTI-S (Figure 2C). Of the 1355 cWindows detected by CACTI-S, 1049 (77.4%) are CACTI-S specific signals. We performed secondary analyses to assess the stability of CACTI-S with respect to window definition. First, we varied the window size from the default 50 kb to 10 kb, 25 kb, and 100 kb. Using the proportion of cPeaks replicated by cWindows as a quantitative measure, we observed highly consistent performance across all window sizes (Supplementary Notes; Figure S1). We further conducted shifting-window analyses by offsetting the window start positions by 25 kb on chromosome 5 in H3K36me3, and found that more than 82.5% of cWindows were replicated before and after shifting (Supplementary Notes; Figure S2). Together, these results demonstrate that CACTI-S is highly robust to alternative window definitions. To further assess the biological validity of CACTI-S signals, we performed annotation- and QTL-based validation analyses. Consistent with the known association of H3K36me3 with transcribed gene regions^23^, CACTI-S cWindows for H3K36me3 showed 3.75-fold enrichment (p-value = 2.47 × 10^-120^) for transcription-associated ChromHMM states^26^ (including Tx5′, Tx, Tx3′, TxWk, TxReg, TxEnh5′, TxEnh3′, and TxEnh). At the QTL level, CACTI-specific cQTLs are significantly enriched in eQTLs from whole blood (OR = 2.07, p-value = 1.86x10^-21^; Figure S3). Moreover, a substantial fraction of H3K36me3 cQTLs overlapped with other transcription-associated QTLs, including 32% overlap with whole blood eQTLs from eQTLGen^25^, 20% with H3K4me1 cQTLs, 27% with H3K4me3 cQTLs in LCLs, and 31% with H3K27ac cQTLs, together explaining the majority (53%) of identified H3K36me3 cQTLs (Methods). Collectively, these quantitative validations indicate that CACTI-S identifies biologically coherent broad regulatory regions and high-confidence cQTLs aligned with established transcriptional regulatory landscapes.

CACTI-S attains substantially higher power than the standard single-peak-based method, not only because multivariate associations are generally more powerful, but also because they overcome the difficulty of accurate peak calling for broad-peak signatures. For example, CACTI-S successfully identifies a significant cQTL signal (lead cQTL SNP p-value = 2.68 × 10^-18^) in a window on chromosome 3 that contains no identifiable peaks and is therefore excluded by the standard peak-based method (Figure 2D). When the peaks of H3K36me3 are defined, CACTI-S still attains higher statistical power with the multivariate approach than the standard univariate association method (an example in Figure S6). We note that for histone marks with narrow peak signatures, such as H3K27ac, peak calling followed by CACTI should be the preferred choice as it outperforms CACTI-S (Supplementary Notes; Figure S7). In summary, CACTI-S significantly outperforms single-peak cQTL mapping for histone marks with broad peak signatures, making it a powerful tool for studying the genetic control of these marks.

### A comprehensive cQTL map across multiple marks and cell types

We applied CACTI and CACTI-S to datasets spanning various histone marks, cell types, and tissues to obtain a comprehensive map of cQTLs. The datasets include ChIP-seq data of H3K27ac, H3K4me1, H3K4me3 in LCL^9^ (N=78, Figure 2E), CUT&TAG data of H3K36me3 in LCL^23^ (N=95, Figure 2E), and ChIP-seq data of H3K27ac, H3K27me3, H3K4me1, and H3K4me3 in uninfected and infected macrophages^14^ (N=35, Figure 2E, Figure S8). We used CACTI to analyze marks with narrow peak signatures (i.e., H3K27ac, H3K4me1, and H3K4me3) and CACTI-S for marks with broad peak signatures (i.e., H3K36me3 and H3K27me3). Consistent with previous findings^14^, we found that cQTLs for the marks H3K4me3, H3K4me1, and H3K27ac are highly shared (Figure S9). Across all datasets, CACTI and CACTI-S consistently outperform the single-peak–based method, identifying 51%–255% more cQTLs (Figure 2E, Figure S10). We observed significant eQTL enrichment among the CACTI- or CACTI-S–specific cQTLs (OR range = 1.03–2.07, p-value range = 4.11x10^-58^–2.48x10^-1^; Figure S2). CACTI and CACTI-S also replicate the majority of cQTLs (98%–100%) detected by the standard mapping methods (Figure S10). Overall, CACTI-specific cQTLs show weaker enrichment than shared cQTLs (Figure S3). This is expected and consistent with CACTI’s increased power to detect weaker regulatory signals that do not reach significance in single-peak analyses and therefore tend to have smaller downstream effects on gene expression.

We also applied CACTI to ChIP-seq data of H3K27ac in brain, heart, lung, and muscle from eGTEx^10^ (N=66–113). The number of H3K27ac cQTLs in eGTEx is significantly lower than in other datasets. For example, we detected only 507 cWindows in the eGTEx brain (N = 113) versus 7,027 cWindows in LCL (N = 78) and 3,166 cWindows in macrophage (N = 35, Figure 2E). We note that while the number of cQTLs we report is much lower than the original eGTEx study^10^, this is explained by the use of a loose p-value threshold of 0.005 and the lack of multiple testing correction in the original study (Figure 2, Figure S11). To reduce false discoveries and ensure a more accurate and fair comparison, we applied multiple testing corrections at 5% FDR to the eGTEx results. The correction reduces the cQTL signals of single peaks by an average of 83% (Figure 2F, Figure S11). In contrast to the significantly larger number of cQTLs detected in other histone mark datasets, CACTI detects comparable numbers of signals to the single-peak-based method in eGTEx tissues under the same FDR threshold (Figure 2E). This diminished performance is likely attributable to a lower effective signal-to-noise ratio in the dataset (Discussion). We therefore focused on LCL and macrophage cQTLs in follow-up analyses.

### CACTI cQTLs explain more GWAS loci than standard cQTLs

To explore the regulatory effects of genetic variants associated with complex traits, we performed colocalization analysis for GWAS loci of 36 blood traits^27^ and 8 immune diseases^28–34^ (44 traits in total) with the comprehensive map of cQTLs across four epigenetic chromatin marks in infected and uninfected macrophages and four marks in LCL (Figure 3A,B, Methods).

**Figure 3.**
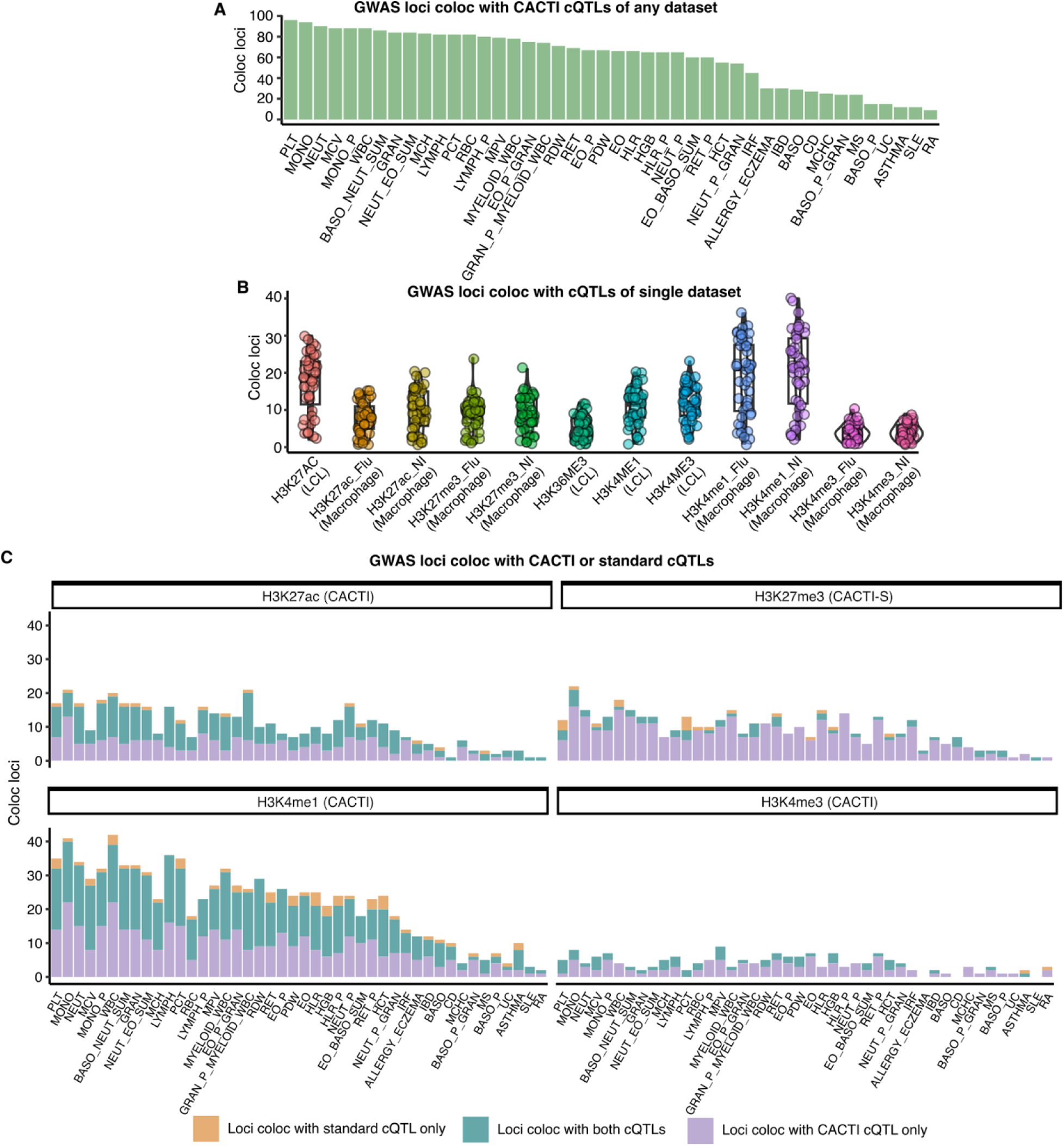
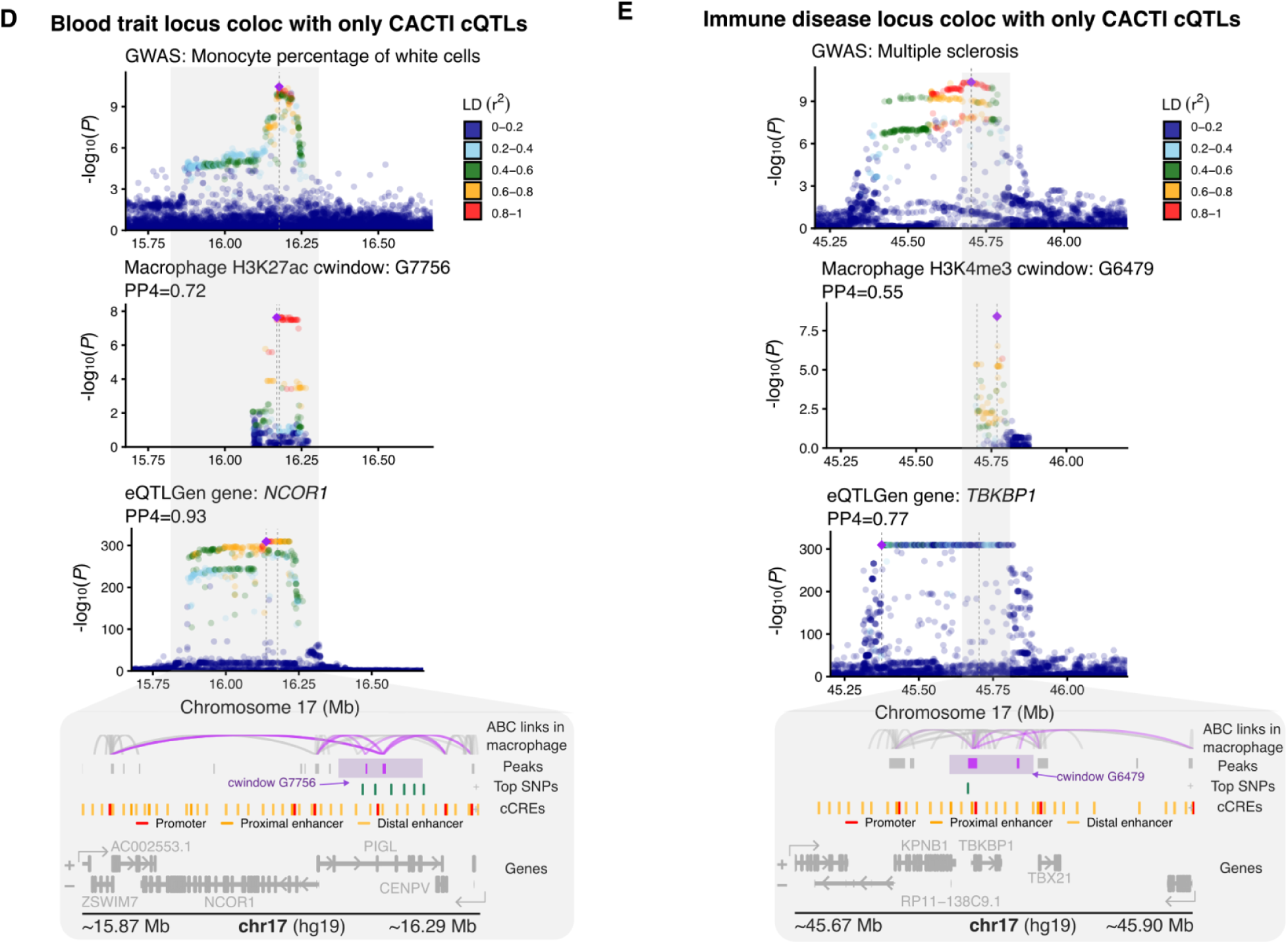
Colocalization of GWAS loci of 36 blood-related traits and 8 immune diseases with cQTLs across marks and cell types. (A) GWAS loci coloc with cQTLs detected by CACTI and CACTI-S. X-axis shows 36 blood related traits and 8 immune diseases used for colocalization with cQTLs. Y-axis shows the number of GWAS loci coloc with cQTLs of any mark in any cell type. Traits are ordered by the total number of colocalized loci. (B) GWAS loci coloc with cQTLs in single datasets detected by CACTI and CACTI-S. X-axis shows each individual dataset of a mark and a cell type. Y-axis shows the number of GWAS loci coloc with cQTLs in the corresponding dataset. Each point represents a trait. (C) GWAS loci colocalize with cQTLs detected by CACTI and CACTI-S and standard mapping methods in uninfected macrophages. The X-axis shows 36 blood-related traits and 8 immune diseases used for colocalization with cQTLs. The cQTLs are from uninfected macrophages. H3K27ac, H3K4me1 and H3K3me4 cQTLs were identified by CACTI, and H3K27me3 cQTLs were identified by CACTI-S. The Y-axis shows the number of coloc loci. Purple bars show the loci colocalized with only CACTI (or CACTI-S) cQTLs. Yellow bars show the loci that colocalize with only standard single-peak–based cQTLs. Blue bars show the loci colocalized with both types of cQTLs. Traits are in the same order as (A). (D) Colocalization between a GWAS locus of monocyte percentage and CACTI cQTLs of H3K27ac in uninfected macrophages. The colocalization of the GWAS locus with eQTLs of *NCOR1* is also shown. Colors represent LD between SNPs and the lead SNP. Bottom genome browser tracks show ABC links in macrophage (highlighting links overlap with peaks included in the cWindow), peaks (highlighting peaks included in the cWindow), cQTL SNPs with top cWindow associations (p-value<10^-6^), candidate cCREs from ENCODE (red promoter, orange proximal enhancer, yellow distal enhancer), and genes around the locus. *NCOR1* has multiple annotated promoters (red bars in the cCRE track). Only one (to the left) is used in the computation of the publicly available ABC scores, which is low due to the lack of contact. The promoter on the right has clear contact with the peaks in the cWindow. (E) Colocalization between a GWAS locus of multiple sclerosis and CACTI cQTLs of H3K4me3 in uninfected macrophages. The colocalization of the GWAS locus with eQTLs of *TBKBP1* is also shown. Colors represent LD between SNPs and the lead SNP. Bottom genome browser tracks show ABC links in macrophage (highlighting links overlap with peaks included in the cWindow), peaks (highlighting peaks included in the cWindow), cQTL SNPs with top cWindow associations (p-value<10^-6^), candidate cCREs from ENCODE (red promoter, orange proximal enhancer, yellow distal enhancer), and genes around the locus.

We grouped GWAS loci into 1 Mb regions, resulting in 361 GWAS loci per trait on average to perform colocalization analysis (Methods, Figure S12). We found that 9 to 96 GWAS loci per trait colocalize with cQTLs of any mark or cell type (defined as a posterior probability of a shared causal variant, PP4>0.5)^10^, corresponding to 6% to 47% of total GWAS loci (mean = 18%; Figure 3A). The proportion of GWAS loci explained by cQTLs is comparable to that explained by eQTLs^14,23^, despite the much smaller sample size of histone mark datasets compared to gene expression datasets. Among cQTLs across marks and cell types, H3K4me1 in uninfected or infected macrophages colocalizes with the largest proportion of GWAS loci (2–40 loci across traits, 1.36%–17.02% of total loci), followed by H3K27ac in LCLs (2–30 loci across traits, 1.27%–10% of total loci) and H3K4me3 in LCL (1–24 loci across traits, 1.92%–10% of total loci). The cQTLs of the repressive mark H3K27me3 colocalize with 1 to 24 (0.77%–7.5%) loci per trait (Figure 3B, Figure S12). H3K36me3 is associated with transcriptionally active genes, and its cQTLs colocalize with 1 to 13 (0.22%–7.02%) GWAS loci per trait (Figure 3B, Figure S12). Over half of the GWAS loci colocalize with more than one mark (Figure S13), suggesting that considering multiple marks offers complementary insights into the mechanisms of GWAS loci.

We next compared cQTL signals identified by CACTI to those from standard single-peak–based mapping methods in terms of their ability to explain GWAS loci. In every case, CACTI-derived cQTLs colocalized with a substantially greater proportion of GWAS hits—an average of 79% more, 94% more, and 182% more for cQTLs of H3K4me1, H3K27ac, and H3K4me3 (respectively) in infected and uninfected macrophages (Figure 3C). Similar proportions were observed for cQTLs in LCL (Figure S12). The difference was even more dramatic for H3K27me3, with nearly all GWAS loci colocalization (69%-100%) only explained by CACTI-S, highlighting the power of this method for capturing genetic associations of chromatin marks that resist peak calling. Most cQTLs identified by standard mapping methods in these datasets were also detected by CACTI; for example, an average of 88% of single-peak H3K4me1 cQTLs in infected and uninfected macrophages also appeared in CACTI.

Overall, CACTI significantly enhances cQTL detection and GWAS colocalization compared to the standard single-peak-based method. This improvement facilitates the functional interpretation of GWAS loci and helps prioritize putative regulatory regions. For instance, CACTI identified cQTLs for a region containing two H3K27ac peaks in macrophages that colocalize with a locus associated with monocyte percentage (PP4 = 0.72; Figure 3D). In contrast, the univariate cQTLs for these two peaks do not exhibit colocalization with a GWAS locus (PP4 = 0.46 and 0.077; Figure S14). Additionally, we detected colocalization (PP4 > 0.5) between this GWAS locus and eQTLs for three genes (*NCOR1*, *CENPV* and *ZSWIM7*, Methods), nominating them as potential *cis* regulatory targets of H3K27ac-marked enhancers (Figure S14). Supporting this, our analysis of Activity-by-Contact (ABC) maps^35^ from macrophage-relevant cell types (Figure 3D; Table S4) revealed that one of the top cQTL SNPs (rs3785625; CACTI p-value=3.2x10^-8^; GWAS p-value=1.3x10^-10^) lies within an enhancer that has strong physical contact with the promoters of all three genes. We note that although public ABC annotations assign *NCOR1* a low score due to the use of a single annotated transcription start site (TSS)^35^, *NCOR1* has an alternative TSS whose promoter-proximal region exhibits clear contact with the peaks in the cWindow (Figure 3D). *NCOR1* (PP4=0.93) is a key regulator of macrophage-mediated inflammation and immune responses^36^, further supporting its potential role in monocyte biology. CACTI thus offers one potential mechanism through which this GWAS locus could operate—namely, altered chromatin conformation at the enhancer in question—that would be overlooked by standard cQTL mapping. In another example, we identified a multiple sclerosis (MS)-associated locus that colocalizes with CACTI-nominated H3K4me3 cQTLs in uninfected macrophages (PP4=0.55; lead p-value=3.9x10^-9^; Figure 3E). The corresponding cWindow contains two distinct peaks, neither of which show significant colocalization with the GWAS locus (PP4 = 0.35 and 0.078; Figure S15) based on their univariate signal strength. The same GWAS locus also colocalizes with eQTLs of three genes, including *TBKBP1*, *NPEPPS*, and *RP11-580I16.2* (Figure S15). Significantly, we found that one of the cQTL peaks is located at the promoter of *TBKBP1* (PP4 = 0.77, Figure 3E), suggesting that the GWAS locus influences *TBKBP1* promoter activity. This positions *TBKBP1*, a gene involved in TNF-α/NF-κB signaling and previously linked to MS, as a strong functional candidate mediating the GWAS association^37–39^. While these two examples will require further functional validation, they demonstrate the utility of CACTI’s increased detection power for nominating plausible regulatory elements that are not visible to existing cQTL mapping techniques, helping to elucidate the functional links between GWAS loci and disease biology.

### CACTI cQTLs provide functional interpretation of GWAS loci beyond eQTLs

Because genetic effects on chromatin configuration may influence gene expression only in specific biological contexts, we expect that many functionally significant regulatory elements will be detectable as cQTLs but not as steady-state eQTLs. To explore this possibility, we investigated how many GWAS loci colocalize with CACTI cQTLs but not eQTLs. Ideally, the eQTLs and cQTLs should be from the same cell type for a fair comparison. However, gene expression datasets of macrophages or LCLs typically have small sample sizes, which could limit the power of colocalization. We found that eQTLs from substantially larger gene expression datasets, such as eQTLGen^25^ (N>30,000), can better recover colocalization signals, even if cell types are not perfectly matched (Figure 4A, Supplementary Notes). We demonstrated this through a comparative colocalization analysis between cQTLs in LCL, eQTLs of GTEx LCL^40^ (N=117) and eQTLs of eQTLGen (whole blood)^25^. For all GWAS loci that colocalize with LCL cQTLs, very few colocalize with only LCL eQTLs (Figure 4A). In contrast, eQTLs from eQTLGen recover most loci that colocalize with LCL eQTLs. We therefore used eQTLGen eQTLs to colocalize with trait and disease loci.

**Figure 4.**
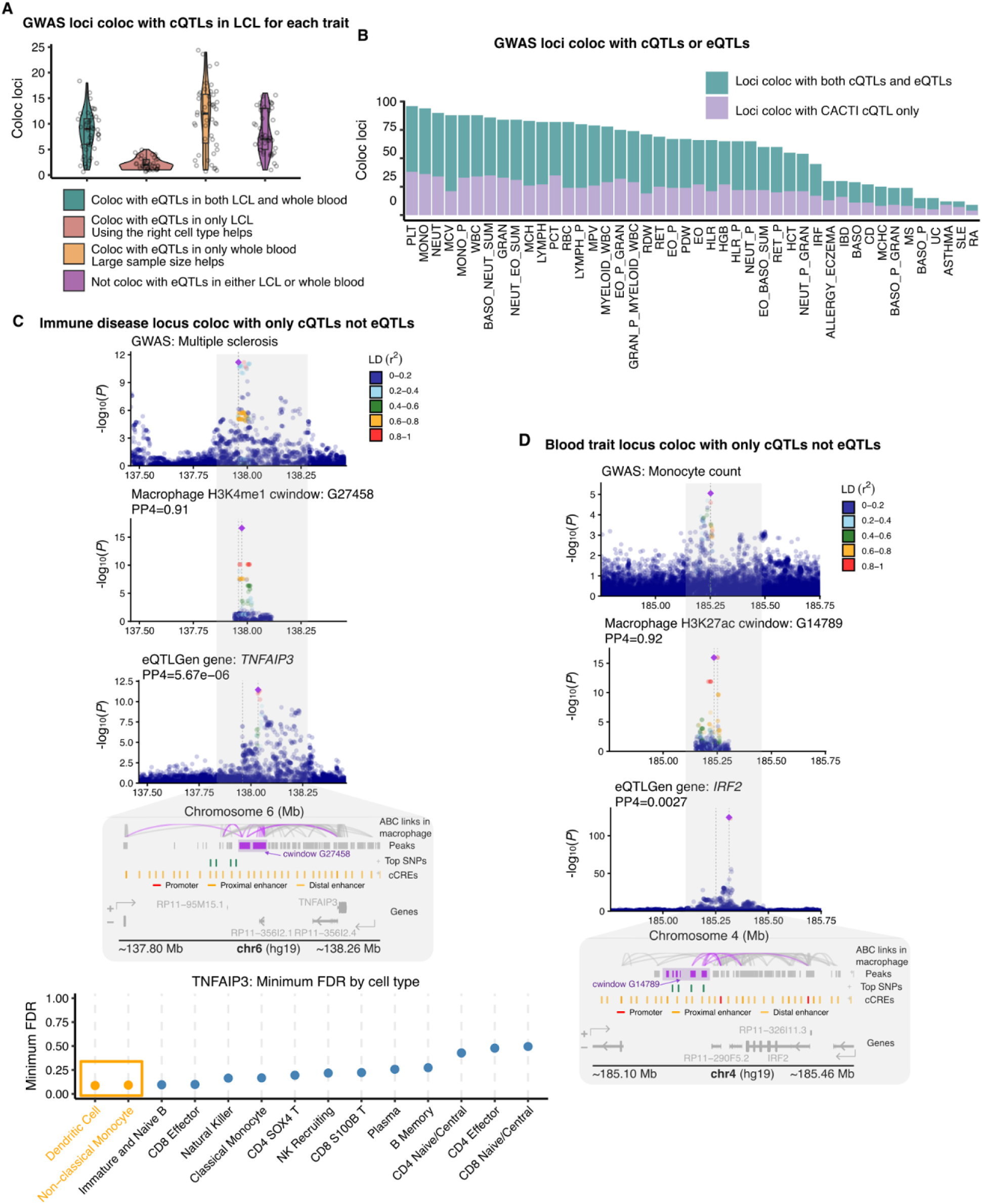
Colocalization of GWAS loci of 36 blood related traits and 8 immune diseases with cQTLs versus eQTLs. (A) GWAS loci colocalize with cQTLs in LCL and eQTLs from LCL or whole blood. Each point is a blood trait or immune disease. Y-axis shows the number of loci colocalize with cQTLs. These loci are categorized into four groups, including loci also colocalize with eQTLs in (1) both whole blood and LCL, (2) only whole blood, (3) only LCL, (4) neither whole blood nor LCL. eQTLs in whole blood are from eQTLGen. eQTLs in LCL are from GTEx. (B) GWAS loci colocalize with only CACTI cQTLs but not eQTLs (purple) or with both cQTLs and eQTLs (blue). eQTLs are from eQTLGen. (C) A GWAS locus of multiple sclerosis on chromosome 6 colocalizes with cQTLs of H3K4me1 in uninfected macrophages identified by CACTI. This locus does not colocalize with eQTLs of any nearby genes, including *TNFAIP3*. Colors represent LD between SNPs and the lead SNP. Bottom genome browser tracks show ABC links in macrophage (highlighting links overlap with peaks included in the cWindow), peaks (highlighting peaks included in the cWindow), SNPs with top cWindow associations (p-value<10^-6^), candidate cCREs from ENCODE (red promoter, orange proximal enhancer, yellow distal enhancer), and genes around the locus, followed by a scatter plot showing the minimum FDR of cell-type–specific eQTLs for *TNFAIP3* across 14 immune cell types in the OneK1K dataset. (D) A GWAS locus of monocyte count on chromosome 4 colocalizes with cQTLs of H3K27ac in uninfected macrophages identified by CACTI. This locus does not colocalize with eQTLs of any nearby genes, including *IRF2*. Colors represent LD between SNPs and the lead SNP. Bottom genome browser tracks show ABC links in macrophage (highlighting links overlap with peaks included in the cWindow), peaks (highlighting peaks included in the cWindow), SNPs with top cWindow associations (p-value<10^-6^), candidate cCREs from ENCODE (red promoter, orange proximal enhancer, yellow distal enhancer), and genes around the locus. See Figure S17 for the scatter plot showing the minimum FDR of cell-type–specific eQTLs for *IRF2* across 14 immune cell types in the OneK1K dataset.

Among all GWAS loci that colocalize with cQTLs of any mark in any cell type, 24% to 75% (4 to 38 loci) did not colocalize with eQTLs of any genes in eQTLGen (Figure 4B, Figure S16), despite the much larger sample sizes of the eQTL dataset. As an illustrative example, a multiple sclerosis (MS) GWAS locus on chromosome 6 colocalizes with H3K4me1 cQTLs (PP4 = 0.91; Figure 4C) but not with any detectable eQTLs. Activity-by-contact (ABC) interaction maps indicate that enhancers within the corresponding cWindow physically interact with the promoter of *TNFAIP3* in macrophages (Figure 4C). *TNFAIP3* is a key negative regulator of inflammation, and reduced *TNFAIP3* expression has been implicated in heightened inflammatory responses and MS pathogenesis^40^. This observation supports the notion that cQTLs can reveal highly cell-type– and context-specific regulatory mechanisms that are not captured by bulk eQTL analyses. To further support this interpretation, we examined *TNFAIP3* eQTLs across peripheral blood mononuclear cell types from the OneK1K dataset^41^. *TNFAIP3* exhibits pronounced cell-type–specific eQTL effects (Figure 4C), with the strongest signals (FDR < 0.1) observed in dendritic cells and non-classical monocytes, which are closely related to macrophages. Together, these results suggest that the regulatory effects captured by cQTLs are active in macrophage and macrophage-like cell types, while steady-state gene expression measured in whole blood lacks the resolution to detect such context-specific effects. In another example, a GWAS locus for monocyte count on chromosome 4 strongly colocalizes with an uninfected macrophages H3K27ac cQTL (PP4=0.92; Figure 4D). While this locus shows no significant colocalization with any genes’ eQTLs (for example coloc PP4=0.0027 between eQTLs of *IRF2* and the GWAS locus), ABC interaction maps indicate that active enhancers in the cWindow physically loop to the promoter of *IRF2* in macrophages (Figure 4D). This is notable, as *IRF2* is known to be essential for the differentiation of non-classical monocytes^42^. Nonetheless, eQTL effects were not observed in macrophage-related cell types in OneK1K (Figure S17). The lack of a co-localizing eQTL signal suggests the genetic effect may operate through enhancer-mediated regulatory mechanisms in a macrophage-specific manner, while the colocalization with our cQTL hints at a possible mechanism involving histone modification. The above examples suggest that many disease-associated variants may exert genetic influences of this kind, altering gene expression only under particular stimuli, in specific cell types, or during distinct cellular states. While such regulation may not be captured by *cis*-eQTL mapping in bulk cell populations, cQTLs, which reflect the dynamic and poised regulatory landscape, appear more robust for detecting these genetic effects across related cell types and states—a finding consistent with both our own expectations and recent literature^15^. In summary, CACTI and CACTI-S cQTLs improve the interpretability of GWAS loci by increasing the number of colocalized loci and reducing the sensitivity to cellular context, highlighting the importance of considering molecular traits beyond gene expression.

## DISCUSSION

We introduced CACTI, a powerful method for identifying cQTLs of multiple nearby peaks from assays of chromatin modification or accessibility using multivariate association tests. CACTI and CACTI-S consistently outperform the standard single–peak–based method by 51%–255% across all histone marks in different cell types (Figure 2E). Using CACTI, we constructed a comprehensive map of cQTLs associated with five epigenetic marks across cell types. CACTI cQTLs can explain between 9 and 96 GWAS loci per trait from a set of 44 complex traits, corresponding to 6% to 47% of total GWAS loci (Figure 3A and 3B). This is comparable to the proportion of GWAS loci explained by eQTLs, even though cQTLs are mapped in a significantly smaller sample size than eQTLs. Furthermore, among all GWAS loci that colocalize with cQTLs, 24% to 75% did not colocalize with eQTLs of any genes (Figure 4B), despite the much larger sample sizes of eQTLs. The priming effects of epigenetic changes can explain some of these signals: genetic effects on epigenetic marks can be observed even when their downstream influence on gene expression cannot be captured by steady-state gene expression or gene expression in the wrong cell type. These findings support the importance of assaying histone modifications and chromatin accessibility alongside gene expression data, as well as the value of CACTI in improving the interpretation of GWAS loci.

CACTI-S skips peak calling and significantly simplifies cQTL mapping for histone marks with broad peak signatures and low signal-to-noise ratios. The advantage of CACTI-S is especially evident when explaining GWAS loci with cQTLs. For example, almost all H3K27me3 cQTLs that colocalize with GWAS loci are mapped by CACTI-S but not by standard single-peak cQTL mapping methods (Figure 3C), presumably because the latter rely on peak calling and thus largely fail to register the relatively diffuse signatures of this epigenetic mark. Nonetheless, we note that CACTI is still superior to CACTI-S when mapping cQTLs in histone marks with narrow peak signatures. For example, while CACTI-S outperforms the single-peak-based method for H3K27ac in LCL, CACTI outperforms both methods (Supplementary Notes). Therefore, we recommend peak calling followed by CACTI for histone marks with a narrow peak signature.

While our analyses focus on datasets of various histone marks, CACTI can also be applied to other chromatin-based assays such as ATAC-seq and DNase-seq, which quantify chromatin accessibility rather than histone modifications. By grouping nearby regions within fixed windows, CACTI can be used to identify chromatin accessibility QTLs (caQTLs). However, certain practical considerations may necessitate changes to the protocol. For one, the optimal window size may differ because ATAC-seq peaks are typically narrower than histone mark peaks. Users may need to experiment with different window sizes to determine the best parameters for their data. The sample sizes of these assays are also typically very small, which can limit the detection power of CACTI and other population-based QTL detection methods.

The complete list of cQTLs generated by CACTI provides a valuable resource for further exploration of epigenetic marks and regulatory elements in molecular and complex traits (Table S1, https://doi.org/10.5281/zenodo.18407484). We performed further analyses linking the colocalized peaks to nearby genes, for example, by using ABC scores^43^. Nonetheless, elucidating the detailed molecular mechanisms involved will require further experimental validation.

A natural extension of CACTI is the joint analysis of peaks from multiple chromatin marks, which could further enhance power and biological insight. In this study, CACTI analyzes one chromatin mark at a time because its multivariate tests rely on accurate estimation of the correlation structure among peaks, which is difficult to achieve across marks assayed in separate experiments due to mark-specific noise and batch effects. In addition, different chromatin marks often represent distinct regulatory states, and naive cross-mark peak grouping would reduce interpretability. We therefore focused on single-mark analyses to ensure statistical robustness, while viewing principled multi-mark integration as an important future direction for CACTI.

### Limitations of the study

Our approach has several limitations. First, the optimal window size used in CACTI to group multiple peaks and map their cQTLs could vary for different epigenetic marks. While our analyses demonstrated that the performance of CACTI was not sensitive to the choice of different window sizes, end users may be able to maximize power by experimenting with different window sizes to determine the optimal window size for their analyses. Furthermore, while we used a fixed window in these analyses, the window size can be defined as dynamic *cis* regulatory domains of various sizes^18^. Second, detection of significant cQTLs for a cWindow containing multiple peaks does not necessarily indicate that the cQTL leads to changes in all peaks. The associated SNP may be associated with one or more peaks in the cWindow. Identifying the associated peaks may thus require combining CACTI analysis with univariate association effects. Third, the colocalization signals of cQTL and GWAS loci do not necessarily imply that disease risk is mediated through epigenetic changes. As with eQTL-GWAS colocalization, these signals should be interpreted with caution and supported by complementary data—such as measures of gene expression or chromatin accessibility in relevant tissues or cell types—and, where possible, by experimental validations. Lastly, while CACTI improves GWAS interpretability by identifying colocalizations with cQTLs that may extend beyond the specific cell type or context analyzed, this strength also represents a potential limitation. By capturing shared genetic effects across related contexts, CACTI may identify associations that are biologically more relevant in other cell types than in the analyzed samples. Such cross-context signals can provide insight into the broader genetic architecture of complex traits, but they also carry a risk of over-interpretation when viewed as context-specific effects. We therefore emphasize that CACTI-identified signals should be interpreted cautiously and, where possible, supported by complementary evidence such as gene expression data, chromatin accessibility assays, or functional annotations.

CACTI’s performance depends critically on data quality and effective signal-to-noise ratio of the data. In the eGTEx datasets, we observed substantially fewer detected cQTL windows despite larger sample sizes. For example, brain tissue with ∼100 samples yielded only ∼400 cQTL windows, whereas a macrophage dataset with 35 samples identified ∼4,000 cQTL windows. This large disparity suggests a lower effective signal-to-noise ratio in the mixed-tissue data, likely limiting the power of multivariate peak modeling. Under such conditions, jointly modeling multiple peaks does not necessarily improve power and may be less effective when correlated signals are weak or inconsistent. Additional factors, including tissue heterogeneity and dilution of cell-type-specific regulatory effects, may also contribute but are difficult to disentangle with the available data.

## STAR METHODS

### RESOURCE AVAILABILITY

#### Lead contact

Further information and requests for resources and reagents should be directed to and will be fulfilled by the lead contact, Xuanyao Liu.

#### Materials availability

No materials were generated in the study.

#### Data and code availability

Publicly available ChIP-seq data for H3K27ac, H3K27me3, H3K4me1, and H3K4me3 in macrophages were obtained from https://zenodo.org/records/10108241. Publicly available H3K27ac ChIP-seq data for brain, heart, lung, and muscle tissues were obtained from the eGTEx v8 project www.gtexportal.org. Whole blood eQTL summary statistics were obtained from eQTLGen https://www.eqtlgen.org/. Publicly available ABC links data were obtained from https://www.engreitzlab.org/resources. GWAS summary statistics for the 36 blood cell traits analyzed in this study are publicly available at https://ftp.sanger.ac.uk/pub/project/humgen/summary_statistics/human/2017-12-12/. GWAS summary statistics for the immune diseases analyzed in this study are publicly available at AE (https://alkesgroup.broadinstitute.org/UKBB/), RA (https://genepi.qimr.edu.au/staff/manuelF/gwas_results/main.html), ASTHMA (https://genepi.qimr.edu.au/staff/manuelF/gwas_results/main.html), SLE (https://www.ebi.ac.uk/gwas/studies/GCST003156), IBD (ftp://ftp.sanger.ac.uk/pub/project/humgen/summary_statistics/human/2016-11-07/), CD (ftp://ftp.sanger.ac.uk/pub/project/humgen/summary_statistics/human/2016-11-07/), UC (ftp://ftp.sanger.ac.uk/pub/project/humgen/summary_statistics/human/2016-11-07/). In addition, all data generated in this study, including the summary statistics of the identified cQTLs and cWindows, are available at https://doi.org/10.5281/zenodo.18407484 or upon request. The R package implementing CACTI and code for major analyses is available on GitHub at https://github.com/XuanyaoLiuLab/cacti.

### METHOD DETAILS

#### CACTI method

CACTI uses a multivariate association test to map cQTLs associated with multiple peaks or segments. CACTI can be applied to both histone marks and chromatin accessibility of narrow peak signatures, such as H3K27ac, H3K4me1, H3K4me3, or ATACseq, and broad signatures, such as H3K36me3 and H3K27me3. For broad peak signatures, we developed segment-based-CACTI (CACTI-S) that skips peak calling and maps cQTLs directly on multiple segments. The details are described below. CACTI (and CACTI-S) has two main steps: 1. Grouping peaks or segments into windows; 2. Multivariate association test to map cQTLs of the windows.

##### 1. Grouping peaks or segments into windows

For histone marks with narrow peak signatures, peaks are first called using MACS2^44^. Then peaks are normalized following the three-step normalization procedure used in Fair et al.^23^: 1) for each peak, we normalized the raw counts into CPM to account for library size; 2) we standardized the peaks by z-score normalization (centering and scaling) across individuals; and 3) we further applied the rank-based inverse normalization to normalize across peaks/segments. This is a form of quantile normalization that ensures different samples have similar distribution.

CACTI uses a fixed-sized window to group peaks. The window size is flexible, and the default window size for CACTI is 50 kb. CACTI groups peaks within consecutive genomic intervals (e.g., 0–50 kb, 50–100 kb), using the peak start sites of the first and last peaks to define the window boundaries. Peaks spanning multiple windows are assigned to the first window they overlap. The windows can be overlapping or non-overlapping as desired by the users. In this paper, to enable meaningful comparison with single-peak–based methods, we used non-overlapping windows, so that each peak is only present in one window and the improved statistical power can be attributed to aggregating correlated peaks with multivariate association tests.

CACTI-S skips peak calling and divides the whole genome into small non-overlapping segments of 5kb. It then quantifies the reads overlapping each segment using featureCounts^45^. Segments that have zero reads in more than 50% samples are then removed, following the procedure used in Aracena et al.^14^. It then computes log2 counts per million (logCPM) of each segment to normalize library size. Additional filtering criteria can be applied to retain higher quality segments. For example, for H3K36me3, which generally represents transcribed regions, we first calculated the proportion of segments overlapping with transcribed regions (for example, 35%). We then ordered the segments by logCPM and only retained the top 35% of the segments for downstream analyses. For the remaining segments, we performed z-score normalization across individuals, and rank-based inverse normalization across segments. We grouped the remaining segments using non-overlapping 50 kb fixed-width windows, similar to the grouping of peaks.

##### 2. Multivariate association test

Step 2 of CACTI involves a multivariate association test assessing the association between each window containing multiple peaks/segments and each SNP in *cis*. The default definition of a *cis-*SNP is one within 100kb of the window, and can be changed as desired. Notably, the multivariate test requires the univariate associations between a SNP and all peaks/segments in the window. Therefore, *cis* variants tested for multivariate association will be limited to the variants shared by all peaks/segments in the window. In the regression model, CACTI uses PCs derived from the peaks and genotypes as covariates to account for global variations (Table S5). We test if a genetic variant is associated with multiple peaks/segments grouped in a same window using the multivariate model as follows,

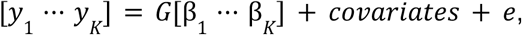

where *G* is the dosage of a reference allele representing the genotype of a SNP, β_*k*_ is the effect of the SNP on *k*-th peak/segment in the window containing *K* peaks/segments, and *y*_*k*_ is the intensity level of the *k*-th peak/segment. To test if a SNP of interest is significantly associated with the window, i.e. any peaks contained in the window, we test the null hypothesis,

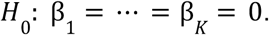

We use a PC-based omnibus test (PCO)^20^, which is a powerful and robust PC-based approach aiming at testing the association with multiple correlated phenotypes with no prior knowledge of the true effects. PCO converts correlated phenotypes into orthogonal PCs. Since multiple PCs are likely to contain association evidence, it is difficult to predict which principal components (PCs) have the largest power to identify shared genetic effects across the phenotypes, PCO utilizes all PCs.

Specifically, PCO uses six different PC-based tests, which combine PCs in linear and non-linear ways, corresponding to different possible architectures of the genetic effects captured on each PC. The test statistic of a single PC is defined as follows:

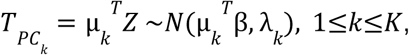

where *Z* is a *K*×1 vector of univariate summary statistic z-scores of the SNP for *K* peaks/segments in a window, µ_*k*_ is the *k*-th eigenvector of the covariance matrix Σ_*K*×*K*_ of *Z*, λ_*k*_ is the corresponding eigenvalue, and β represents the true causal effect. PCO combines six PC-based tests, including,

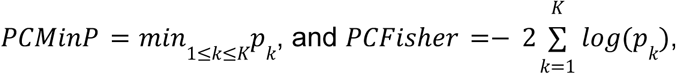

where *p*_*k*_ is the p-value of *T_PC_k__*. These two tests take the best p-value of all single PC-based tests and combine multiple PC p-values into the test statistic. Other tests include,

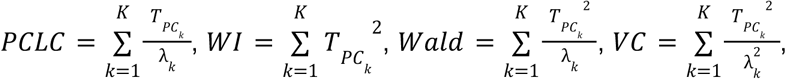

which are linear and quadratic combinations of each single PC-based test weighted by eigenvalues. The six tests achieve best power in specific genetic settings with different true causal effects. PCO takes the best p-value of the PC-based tests as the final test statistic,

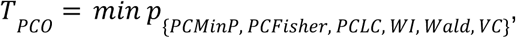

to achieve robustness under unknown genetic architectures while maintaining a high power. The p-value of PCO test statistics can be computed by performing an inverse-normal transformation of the test statistics,

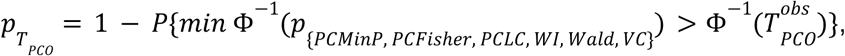

where Φ^−1^ denotes the inverse standard normal cumulative distribution function. The p-value can be efficiently computed using a multivariate normal distribution as described in Liu et al^20^.

For windows containing only one peak/segment, we used the standard univariate association test to obtain p-values. To adjust for multiple testing, we selected the top associated SNP for each window and used Storey’s q-value^22^ to estimate false discovery rate (FDR) across all windows tested, regardless of the number of peaks contained within each window.

### QUANTIFICATION AND STATISTICAL ANALYSIS

#### cQTL mapping of H3K27ac, H3K4me3, H3K4me1, and H3K36me3 in LCL

Univariate cQTL mapping of four histone marks—H3K27ac, H3K4me3, H3K4me1, and H3K36me3—in LCLs was performed by Fair et al.^23,24^ For H3K27ac, H3K4me3, and H3K4me3, peaks were called using MACS2^44^. Raw counts were normalized into logCPM, then standardized by z-score normalization (centering and scaling) across individuals. Finally, rank-based inverse normalization was performed across peaks. For H3K36me3, standard peak calling with MACS2 was not accurate, so the number of reads in transcribed regions of each gene was quantified. Each “peak” is defined as a gene’s transcribed region. logCPM was computed to account for library size. Z-score normalization across individuals and rank-based inverse normalization across peaks was performed. PCs derived from the normalized peaks were used as covariates for QTL mapping (Table S5), which was performed using QTLTools^23,24^. A cis window of 500 kb was used. To claim significant cPeaks, they adjusted p-values of all peaks with Storey’s q-value method^22^ to account for the false discovery rate.

We used CACTI to map cQTLs of H3K27ac, H3K4me3, and H3K4me1. We used non-overlapping windows of 50 kb in the primary analysis, so that each peak is present in only one window. This is to ensure a fair comparison between single-peak–based methods and CACTI. Notably, univariate mapping tested a wider *cis* region compared to our multivariate test, which requires the univariate associations between each variant and all peaks contained within the test window. CACTI decomposed the peaks in each window into PCs and used PCO to compute the associations between each SNP and window. To claim significant cWindows, we selected the top associated SNP for each window and adjusted p-values of all windows with Storey’s q-value method^22^ to account for the false discovery rate.

We used CACTI-S to map cQTLs of H3K36me3. CACTI-S quantified reads in each 5kb consecutive segments with featurecount^45^ and computed logCPM of each segment. Next, we retained high-quality segments for downstream analyses. We found that 35% of segments overlapped with transcribed regions, and therefore, we ranked the segments by logCPM and only retained the top 35% of segments. The retained segments were normalized across samples with z-score normalization and across segments with rank-based inverse normal. We grouped the segments into non-overlapping windows of 50 kb. CACTI-S then uses PCO to test associations between *cis* SNPs and windows. To claim cWindows, we adjusted p-values of all windows with Storey’s q-value method^22^ to account for the false discovery rate.

#### cQTL mapping of H3K27ac, H3K4me3, H3K4me1 and H3K27me3 in infected and non-infected macrophage

Univariate cQTL mapping of the four histone marks H3K27ac, H3K4me3, H3K4me1, and H3K27me3 in uninfected and influenza-infected macrophages was performed by Aracena et al.^14^ Specifically, peaks were called using MACS2^44^, then normalized by library size using TMM and converted to adjusted logCPM^14^. The peaks were further normalized across samples by z-score normalization and normalized across peaks by rank-based inverse normalization. Batch, age, the first genetic PCs, and PCs derived from normalized peaks were used as covariates (Table S5). QTL mapping was performed using R package Matrix eQTL^46^. The *cis* region used to select SNPs for association testing was defined as 100kb.

We used CACTI to map cQTLs of H3K27ac, H3K4me3, and H3K4me1. Non-overlapping 50-kb fix-sized windows were used to group peaks in windows. To claim significant cWindows, we adjusted p-values of all windows with Storey’s q-value method^22^ to account for the false discovery rate.

We used CACTI-S to map cQTLs of H3K27me3. CACTI-S quantified reads in each 5kb consecutive segment with featurecount^45^. Low quality segments with reads in less than 50% samples (14 samples) were removed. The segments were normalized for library size with TMM-adjusted logCPM. They were further normalized across samples with z-score normalization, and across segments with rank-based inverse normalization. We grouped the segments into non-overlapping windows of 50 kb. CACTI-S then uses PCO to test associations between *cis* SNPs and windows. To claim cWindows, we adjusted p-values of all windows with Storey’s q-value method^22^ to account for the false discovery rate.

#### Permutation analyses to assess false positive rates

To evaluate the statistical calibration of CACTI and quantify the type I error (T1E) rate, we conducted a permutation analysis on chromosome 5 of the H3K27ac dataset. We performed 10 independent permutations. In each permutation, we randomly permuted both the sample labels and peak labels in the phenotype matrix. We then performed association testing on these null datasets. The T1E rate was calculated as the proportion of windows identified as significant at nominal p-value thresholds of 0.01 and 0.05. For each threshold, we computed the mean T1E rate and the standard error across the 10 permutations. The observed T1E was compared to the nominal thresholds to verify that the method accurately controls error rates under the null hypothesis.

#### Comparison of signals between marks and conditions

##### Shared and unique cQTLs across infected and uninfected conditions in macrophages

We compared the significant cWindows identified in macrophages under uninfected (NI) and influenza-infected (Flu) conditions. This analysis was conducted for H3K27ac, H3K4me1, and H3K4me3 using the standard CACTI framework, and for H3K27me3 using CACTI-S. We assessed the overlap between conditions by determining if a cWindow identified in the discovery condition (e.g., NI) overlaps with a cWindow identified in the replication condition (e.g., Flu). A cWindow was defined as shared (replicated) if it reached significance (FDR<0.05) in the discovery condition and its corresponding overlapping window in the replication condition also met a significance threshold. In contrast, signals were considered unique to a condition if the discovery cWindow was significant but either lacked an overlapping window in the replication condition or the overlapping window failed to meet the significance threshold. To account for potential weak signals and assess the sensitivity of our replication estimates, we performed this comparison using two significance levels for the replication set: a common threshold (FDR<0.05) and a relaxed threshold (FDR<0.1). The proportion of shared versus unique signals was then quantified across all four marks.

##### Sharing and specificity of cQTLs across marks in LCL and macrophages

To assess the sharing and specificity of cQTLs across different marks, we compared cWindows identified for various marks within the same cell type, LCL or macrophages. This analysis included H3K27ac, H3K4me1, and H3K4me3 (narrow marks) analyzed by CACTI, as well as H3K36me3 and H3K27me3 (broad marks) analyzed by CACTI-S. For each pair of marks, we designated one as the reference and the other as the comparison mark. A cWindow from the reference set was considered shared if it achieved significance at FDR<0.05 and overlapped with at least one cWindow (FDR<0.05) in the comparison mark set. We quantified the sharing proportion as the ratio of replicated cWindows to the total number of cWindows identified for the reference mark.

#### Overlap of H3K36me3 cQTLs with other cQTLs and eQTLs

Significant H3K36me3 cQTLs were defined as lead SNPs of cWindows. H3K36me3 cQTLs were compared to cQTLs of reference marks, including H3K4me1, H3K4me3, and H3K27ac. For each reference mark, SNP-level association signals were collapsed by taking the minimum p-value across all tested windows per SNP. A lead H3K36me3 SNP was considered overlapping if it was also significant in the reference dataset (p-value<10^-4^). H3K36me3 cQTLs were additionally compared with eQTLs from the eQTLGen consortium. Overlap proportions were calculated relative to the total number of significant H3K36me3 cQTLs. In addition to overlap specific to individual reference dataset, an overall overlap with any cQTL or eQTL was also defined.

#### Enrichment analyses of CACTI cQTLs for eQTLs

To assess the functional relevance of the identified cQTLs, we quantified their overlap with eQTLs obtained from the eQTLGen consortium. This analysis was performed for LCLs and macrophages and multiple marks (H3K27ac, H3K4me1, H3K4me3, H3K36me3, and H3K27me3).

For each dataset, tested windows were classified as either significant cQTLs (lead SNPs of cWindows) or non-significant background windows. To specifically evaluate the validity of novel findings, significant cQTLs were further stratified into three categories: “All” (the total set of significant cQTLs), “Shared” (cQTLs that replicated as by the single-peak mapping method), and “New” (cQTLs that were novel and not replicated by the single-peak mapping method). We mapped the lead variants of all tested windows to the eQTLGen summary statistics. Enrichment was assessed using a one-sided Fisher’s exact test by comparing the proportion of variants overlapping significant eQTLGen eQTLs in the cQTL categories versus the non-significant background windows.

As a secondary analysis, we re-evaluated eQTL enrichment using background windows matched on the number of contained peaks to control for potential biases arising from differences in peak density. We performed a matched enrichment analysis using mixed-effects logistic regression. Windows were grouped into matched sets based on the number of peaks they contained, and for each window we selected the top associated SNP. We then tested whether the top SNPs of cWindows were more likely to overlap a eQTL, while accounting for heterogeneous window sizes across matched sets by including a random intercept for each set. Using this approach, we observed that top SNPs of cWindows still had high enrichment, indicating that the identified cWindows are enriched for eQTLs even after controlling for peak density.

#### Colocalization of GWAS loci with cQTLs and eQTLs

We performed colocalization analysis between 44 GWAS loci (36 loci for blood traits^27^ and 8 for immune diseases^28–34)^ and multiple types of molecular QTLs, including CACTI cQTLs of various histone marks in macrophage and LCL, standard peak-based cQTLs in macrophage and LCL, and eQTLs in eQTLGen of whole blood and GTEx LCL.

To define regions for colocalization, we first selected the GWAS lead SNP with the most significant p-value for a given locus and expanded a 500kb flanking genomic region centered at that variant. For each GWAS locus, a colocalization test was performed for all molecular phenotypes, including windows, peaks, or genes, that shared at least 150 overlapping *cis* SNPs with the GWAS variants. For eQTLs, we included all genes with transcription start sites located within 1 Mb of the GWAS lead variant. We used the ‘coloc.abf’ function from coloc package^47^ with the default priors. We considered a locus colocalized when the molecular phenotype displayed a significant signal (cWindow, cPeak, or eGene) and PP4 > 0.5. We used LiftOver^48^ to convert genomic coordinates for GWAS loci to align them with molecular QTLs with different genome assemblies, i.e. reads in LCL datasets were aligned to the hg38 human reference genome, while reads in other datasets were based on genome version hg19. We visualized the colocalized regions using LocusCompareR^49^.

We repeated the colocalization analyses for the remainder of the genome, continuing to search for lead SNPs until no additional significant hits (p-value smaller than 10^-7^) could be found.

## ACKNOWLEDGMENTS

We thank Y. Li, Z. Liu, B. Fair, Z. Mu, C.F. Buen Abad Najar and K. Aracena for helpful discussions. We thank F. Naumann and S. Sumner for editing the manuscript. This work was completed in part with resources provided by the University of Chicago’s Research Computing Center. This research was funded by the NIGMS Maximizing Investigators’ Research Award (R35GM138084).

## AUTHOR CONTRIBUTIONS

L.W. and X.L. developed the method. L.W. implemented the method and performed data analyses under the supervision of X.L.. L.W. and Z. Q. implemented the method in R package. L.W. and X.L. wrote the manuscript.

## SUPPLEMENTAL INFORMATION

### SUPPLEMENTAL NOTES

#### CACTI and CACTI-S detection power is robust to the choice of window size

To evaluate how the choice of window size used to group peaks impacts the power of CACTI and CACTI-S, we experimented with different window sizes, including (1) 10kb, 25kb, and 50 kb for three narrow marks H3K27ac, H3K4me1, and H3K4me3 in across two conditions in macrophage, (2) 10 kb, 25 kb, 50 kb, and 100 kb for the broad mark H3K36me3 in LCL (Figure S1). To evaluate the methods’ performance under different window sizes in a fair way, we used the metric of signal replication, which is the number of cPeaks identified by the single-peak mapping method that were also contained in a cWindow detected by CACTI or CACTI-S. Figure S1 shows that the number of replicated cPeaks or the missed cPeaks stay approximately consistent across different window sizes for all marks (Figure S1). We did not compare the total number of cWindows or the number of new cWindows identified by CACTI or CACTI-S under different window sizes, as these two metrics are not for fair comparison and they can change with the window size not relevant to the CACTI’s power.

#### CACTI and CACTI-S are stable to the window start positions

To define non-overlapping windows, peaks or segments were grouped into fixed-width windows of 50 kb based on their start positions. To evaluate the robustness of these window start positions, we performed a sensitivity analysis by repeating the grouping procedure with a 25kb offset (half-window shifting). We assessed the replication between the original and offset windows: a cWindow identified in the original windows was considered replicated if at least one of its constituent peaks or segments was also captured within a significant window in the offset analysis. This analysis ensures that the reported associations are driven by robust underlying signals rather than artifacts of arbitrary window placement.

#### CACTI outperforms CACTI-S for marks with accurate peak calling

We note that for marks with accurate peak calling (such as H3K27ac), CACTI outperforms CACTI-S (Figure S7). To gain more insights into the performance of CACTI-S, we also applied CACTI-S to the narrow mark H3K27ac from LCL. We compared the identified signals for this mark by CACTI and CACTI-S and examined if the signals detected by the two methods overlap. We found that CACTI has better performance than CACTI-S when being applied to narrow marks (Figure S7). Specifically, 90% of cWindows identified by CACTI-S are replicated by CACTI, whereas 66% of cWindows detected by CACTI do not overlap with any cWindows by CACTI-S, representing the new signals specific only to CACTI (Figure S7). These observations suggest that the performance of CACTI-S depends on the accuracy of peak calling. In the case where peaks are well defined, it gains more power by leveraging correlation among multiple peaks and testing them jointly, than cutting the peak regions into smaller segments, which may disrupt the well-defined peak regions and introduce noise into the test. In contrast, in the case where peak boundaries are poorly defined, cutting the region into finer-scale can decrease the effect of inaccurate peak calling on the association testing and obtain a better power.

#### Signal replicability under random sample splitting

To assess the robustness and replicability of detected signals across independent datasets, we performed a split sample replication analysis. Starting from the full samples of H3K27ac in LCL, we generated independent random splits of 72 samples into two non-overlapping halves of 36 individuals. For each split, all downstream analyses, including phenotype processing, covariate adjustment, genotype subsetting, and association testing, were conducted independently within each subset. For each split, associations were calculated using both CACTI and the single-peak–based method. Replication rate was then evaluated by taking the set of significant associations identified in one split and calculating the proportion that were significant in the other split. We used a replication p-value threshold of 0.05 divided by the total number of signals in the first split. This procedure was repeated for two independent random splits to account for variability due to sampling. Higher replication rates indicate more robustness of detected signals to random sample splitting of individuals.

#### Enrichment of H3K36me3 cWindows in regulatory annotations

H3K36me3 peaks are known to usually correspond to transcribed regions. To evaluate whether CACTI-S cWindows for H3K36me3 correspond to known transcribed regions, we compared them to chromatin 25-state ChromHMM annotations for GM12878 lymphoblastoid cell line. We focused on states associated with transcription activity, including Tx5′, Tx, Tx3′, TxWk, TxReg, TxEnh5′, TxEnh3′, and TxEnhW. We computed the overlaps between H3K36me3 cWindows and transcription-associated ChromHMM states. To assess enrichment, we generated a matched set of random genomic windows, which preserves the number and width distribution of the original windows. Enrichment was quantified using a one-sided Fisher exact test to evaluate whether CACTI-S cWindows were more likely to overlap H3K36me3-associated states compared to random expectation.

### SUPPLEMENTAL FIGURES

**Figure S1.**
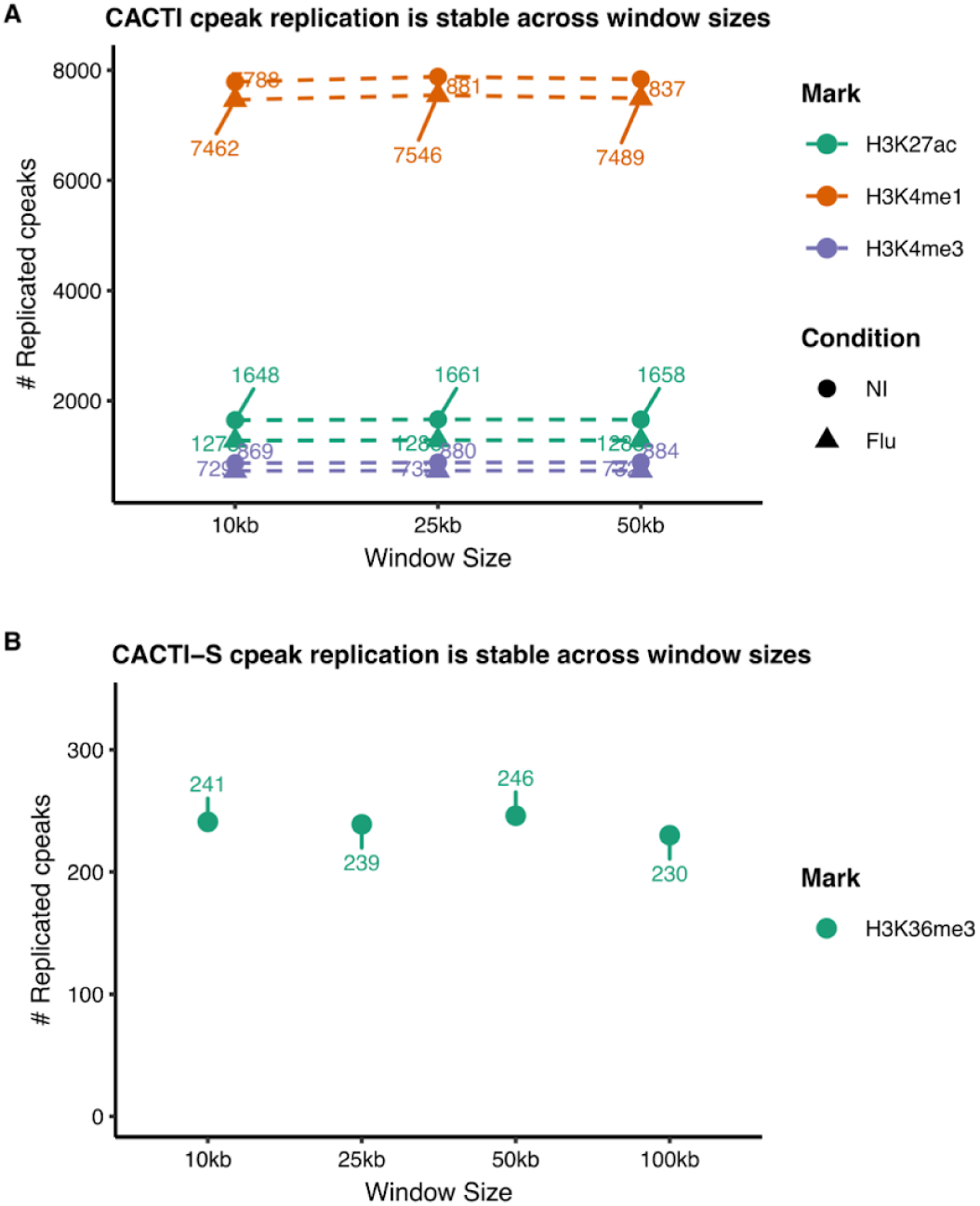
CACTI detection power is robust to the choice of window size. Window sizes used for (A) CACTI and (B) CACTI-S are shown on the x-axis. Results for different marks are shown in colors, conditions are shown in different shapes. Y-axis gives the number of cPeaks that are replicated by CACTI or CACTI-S signals, i.e. these cPeaks are included in a cWindow identified by CACTI or CACTI-S. A larger window, i.e. 100kb, was additionally used for the broad mark H3K36me3.

**Figure S2.**
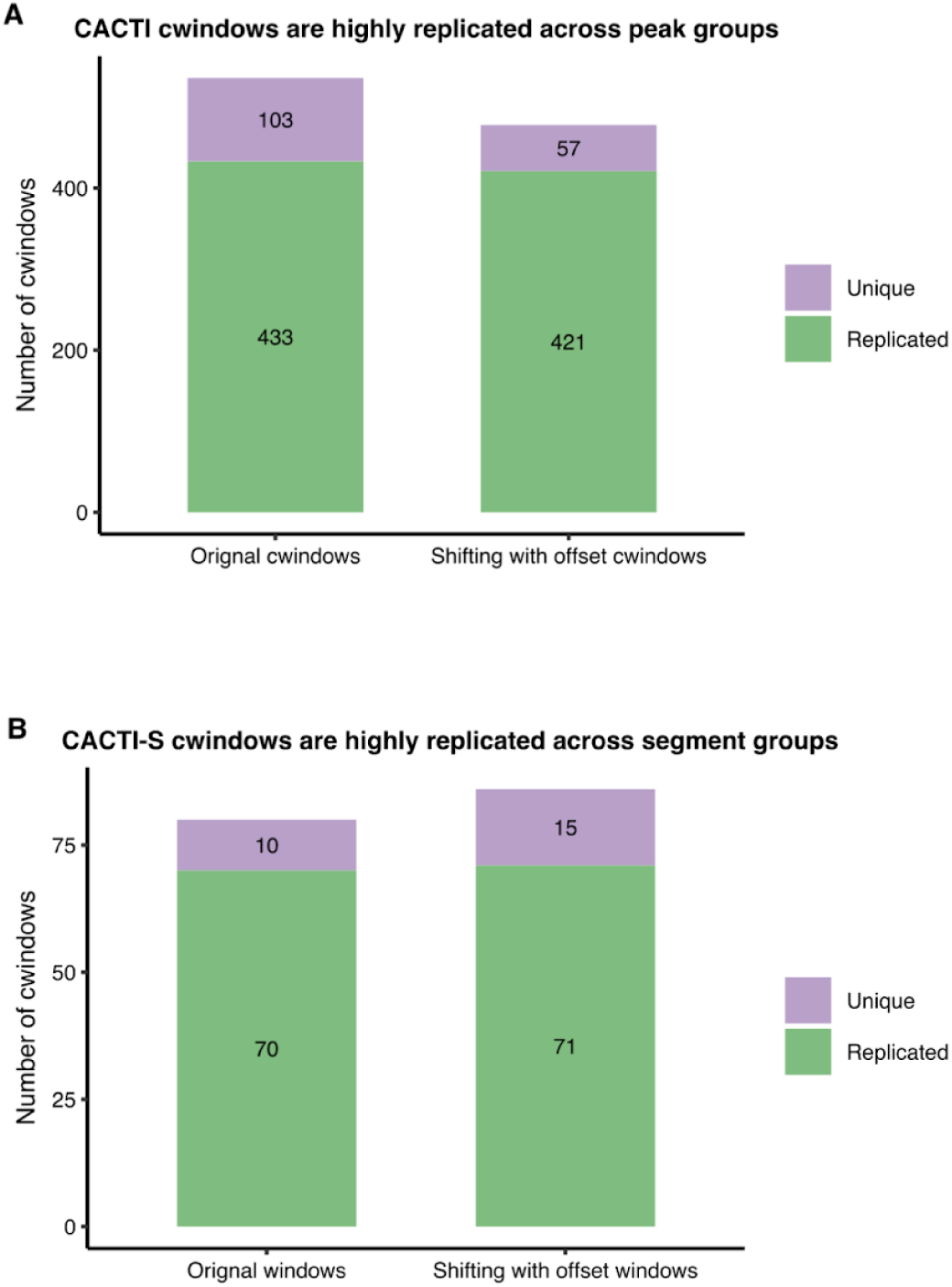
CACTI and CACTI-S are stable to the window start positions. To evaluate the robustness of window definitions, window grouping was performed for (A) H3K27ac peaks and (B) H3K36me3 segments using two ways: the original non-overlapping 50 kb windows and a secondary shifted windows by a 25kb offset. The y-axis displays the number of cWindows identified on chromosome 5 for each mark. A cWindow is classified as “Replicated” if it shares at least one constituent peak or segment with a cWindow in the alternative window definition, demonstrating that the signals are robust to the window start positions. A cWindow is classified as “Unique” if its constituent peaks or segments do not map to a cWindow in the alternative window definition. Numbers inside bars indicate the specific window counts for each category.

**Figure S3.**
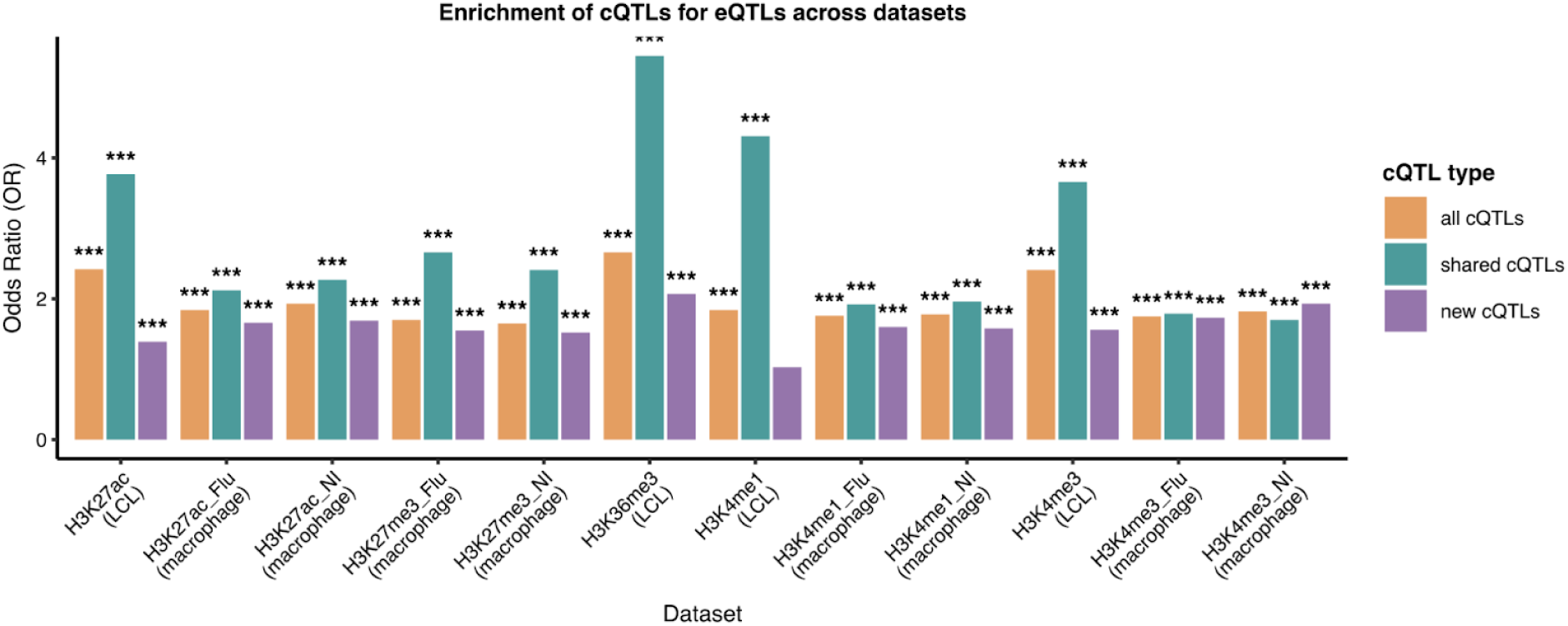
Enrichment of cQTLs detected by CACTI and CACTI-S for eQTLGen eQTLs across marks and cell types. Marks H3K36me3 and H3K27me3 were analyzed by CACTI-S, and the other marks were analyzed by CACTI. The bar plot displays the enrichment odds ratio (OR) from one-sided Fisher’s test. cQTLs are categorized as “All” (detected by CACTI/CACTI-S), “Shared” (detected by both CACTI/CACTI-S and single-peak mapping method), and “New” (detected by only CACTI/CACTI-S but not single-peak mapping method). Asterisks denote statistical significance of the enrichment, * p<0.05, ** p<0.01, *** p<0.001.

**Figure S4.**
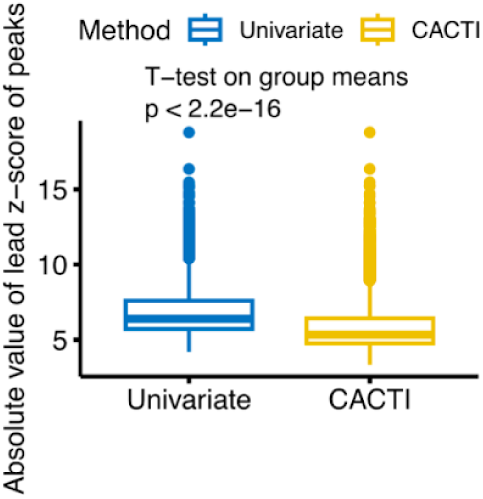
CACTI outperforms the single-peak mapping primarily by capturing signals with weaker effects. Comparison of z-score magnitude of signals by single-peak mapping and CACTI. Univariate represents the single-peak–based mapping method. Y-axis gives the absolute value of z-scores of single peaks. For the univariate method, we focus on the cPeaks and take the p-value of the lead SNP for each cPeak for comparison. For CACTI, we focus on the cWindows and take the minimum p-value of lead SNPs across all peaks contained in the windows. Group means were compared using a two-sample t-test.

**Figure S5.**
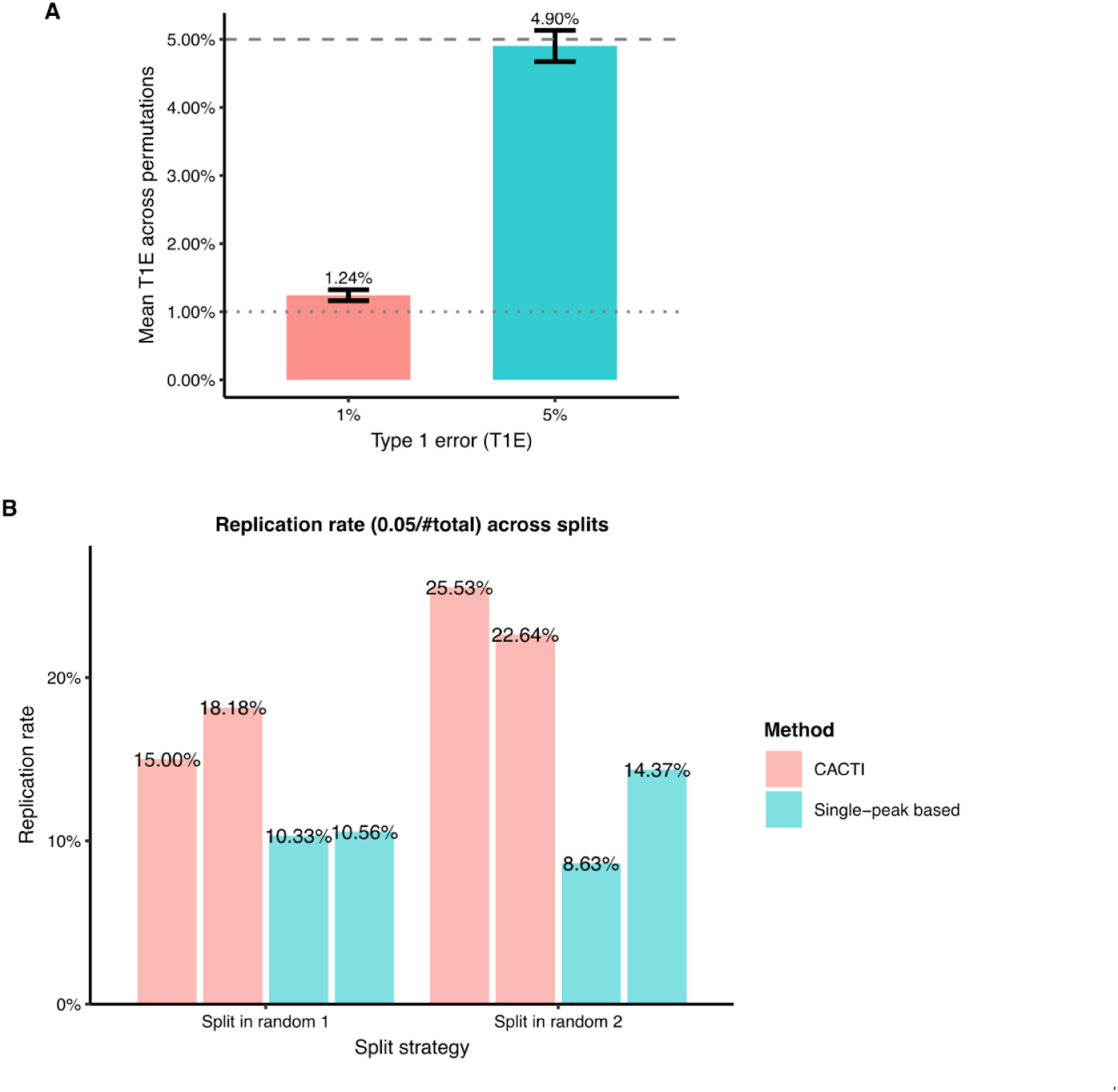
Permutation and replication analysis. (A) T1E for H3K27ac on chromosome 5 in permutation analysis. We performed 10 independent permutations where sample labels and peak labels were shuffled. The bar plot shows the mean observed T1E rate (labelled on the bars) across 10 permutations at two significance thresholds 0.01 and 0.05. Error bars represent the standard error. (B) Signal replicability under random sample splitting. Replication rates of signals identified by CACTI (pink) and the single-peak–based method (blue) under two independent random sample splits (x-axis). We used H3K27ac in LCL of 72 samples. In each split, individuals were randomly partitioned into two subsets (36/36), and signals in one subset were evaluated for replication in the other. Bars show the proportion of signals that replicate across splits, defined as the fraction of signals in one split that also reach significance in the paired split. Percentages above bars indicate replication rates for each method and split. Across both random splits, CACTI consistently achieves higher replication rates than the single-peak–based approach, indicating improved robustness of detected signals under reduced sample size.

**Figure S6.**
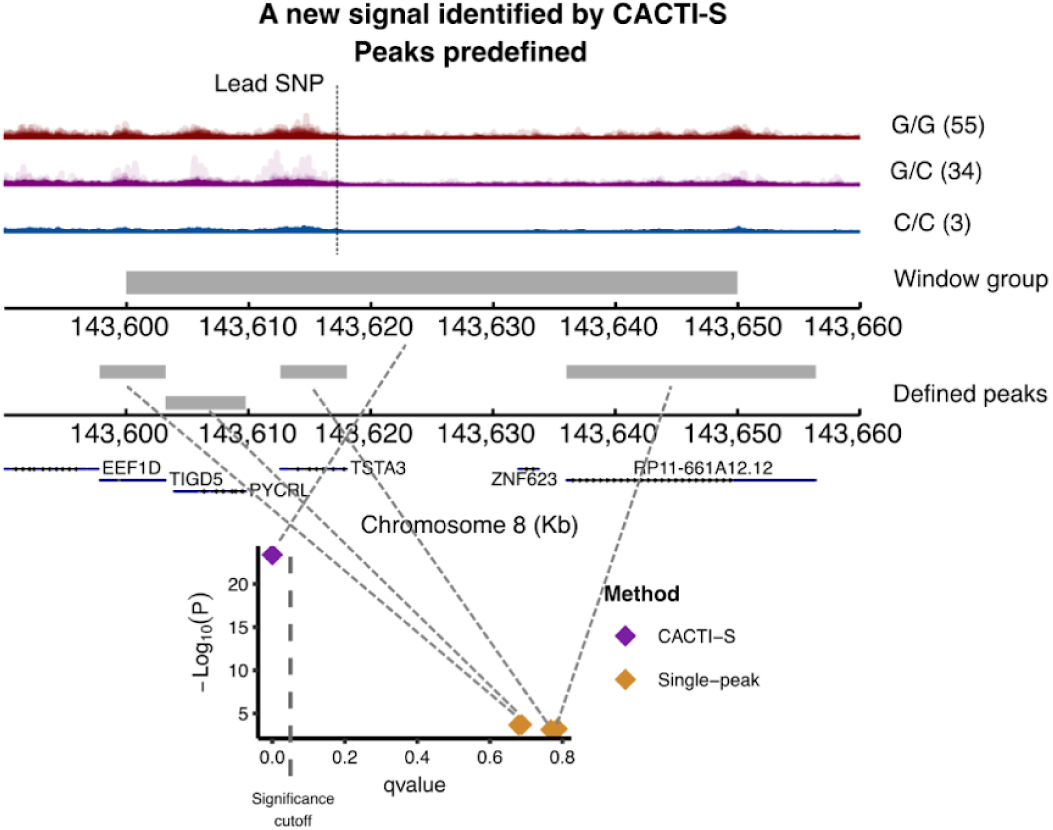
Another example of a new cWindow signal discovered by CACTI-S where peaks were predefined. The upper track displays the aligned reads for individuals classified by the genotypes of the lead cQTL in this region (marked by a dashed line). The middle track represents the cWindow region. The track below shows the peaks predefined in this window region. The bottom track shows the four single peaks overlapped with the window. The scatter plot shows the p-value and q-value from association tests of the window and single peaks.

**Figure S7.**
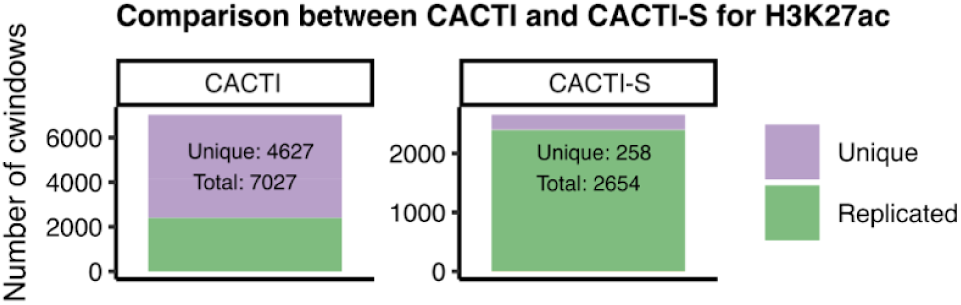
Comparison of signals detected by CACTI and CACTI-S for the mark H3K27ac in LCL. Y-axis represents the number of Unique or Replicated cWindow by CACTI or CACTI-S. A cWindow identified by a reference method (e.g. CACTI) is classified as “Replicated” if it overlaps with a cWindow in the compared method (e.g. CACTI-S). A cWindow is classified as “Unique” if it does not overlap with any cWindow of the other method.

**Figure S8.**
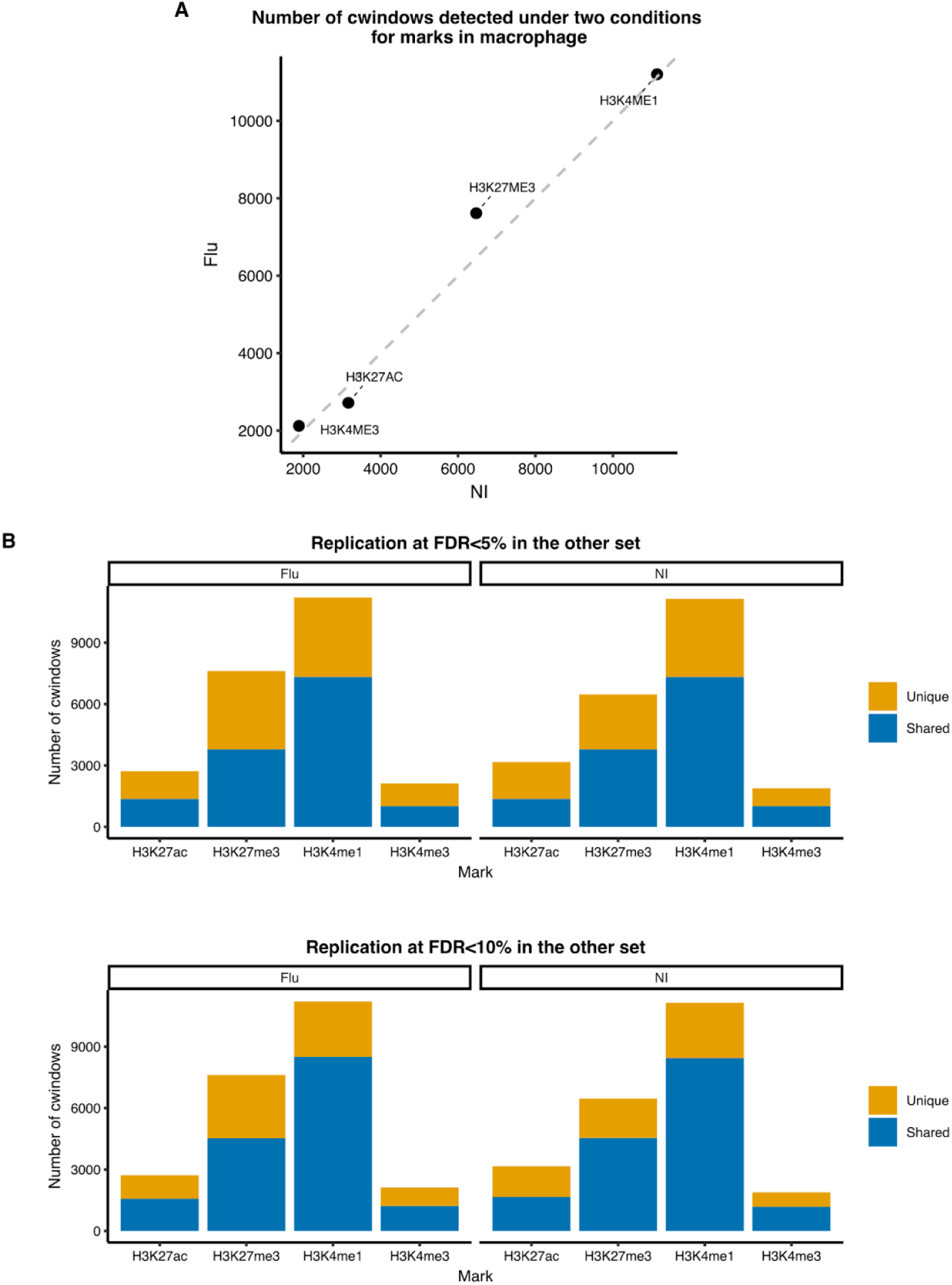
Comparison of cWindows across two conditions in macrophage. (A) Number of cWindows across conditions. X-axis and y-axis represent the number of cWindows identified in the condition of uninfected (NI) and infected (Flu). We used CACTI to analyze marks with narrow peak signatures (i.e., H3K27ac, H3K4me1 and H3K4me3) and CACTI-S for marks with broad peak signatures (i.e. H3K27me3). (B) Replication of cWindows between uninfected (NI) and infected (Flu) conditions. Bar plots illustrate the proportion of cWindows that are Shared or Unique to a specific condition for each mark. A cWindow is defined as “Shared” if it is significant in the discovery condition (FDR<0.05) and overlaps with a cWindow in the replication condition. Two FDR levels for replication are shown: the top panel requires a replication significance of FDR<0.05, while the bottom panel uses a threshold of FDR<0.1.

**Figure S9.**
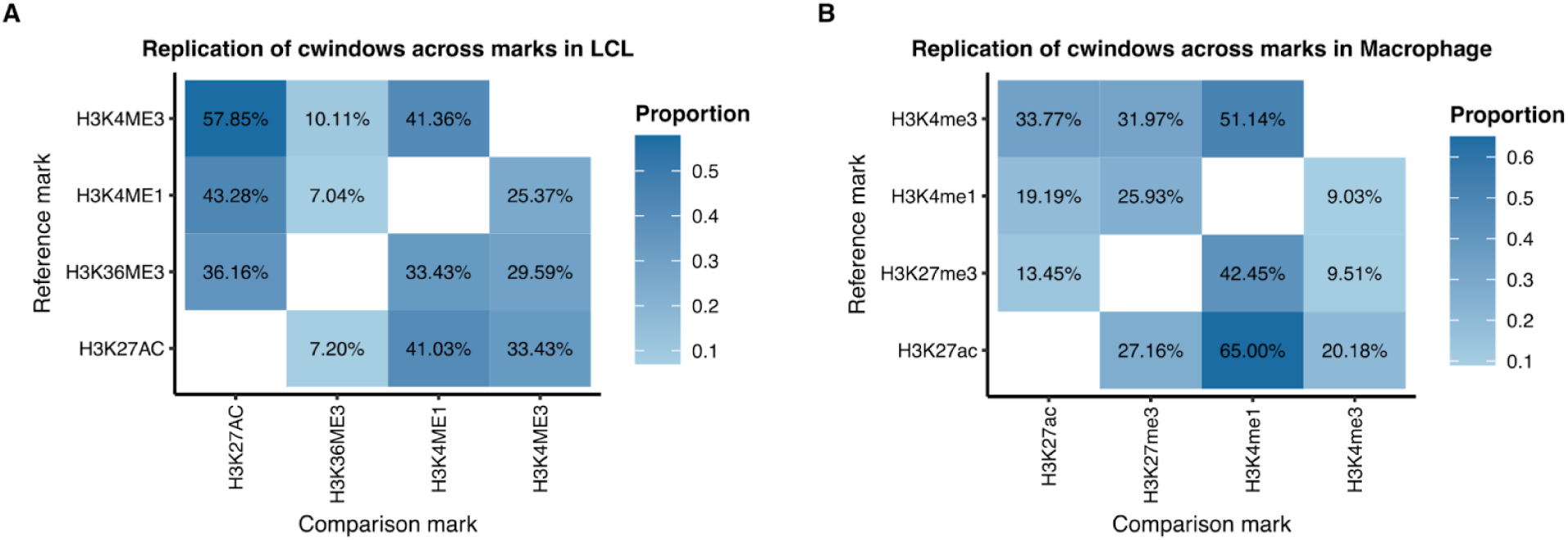
Sharing and specificity of cQTLs across marks. Heatmaps illustrate the proportion of shared cQTL signals between pairs of marks in (A) LCL and (B) uninfected macrophages. Each tile represents the replication proportion, defined as the percentage of significant cWindows (FDR<0.05) for a reference mark (y-axis) that physically overlap with a cWindow for a comparison mark (x-axis). Sharing was assessed for marks with narrow peak signatures (H3K27ac, H3K4me1, and H3K4me3) using CACTI, and for marks with broad signatures (H3K36me3 in LCLs and H3K27me3 in macrophages) using CACTI-S.

**Figure S10.**
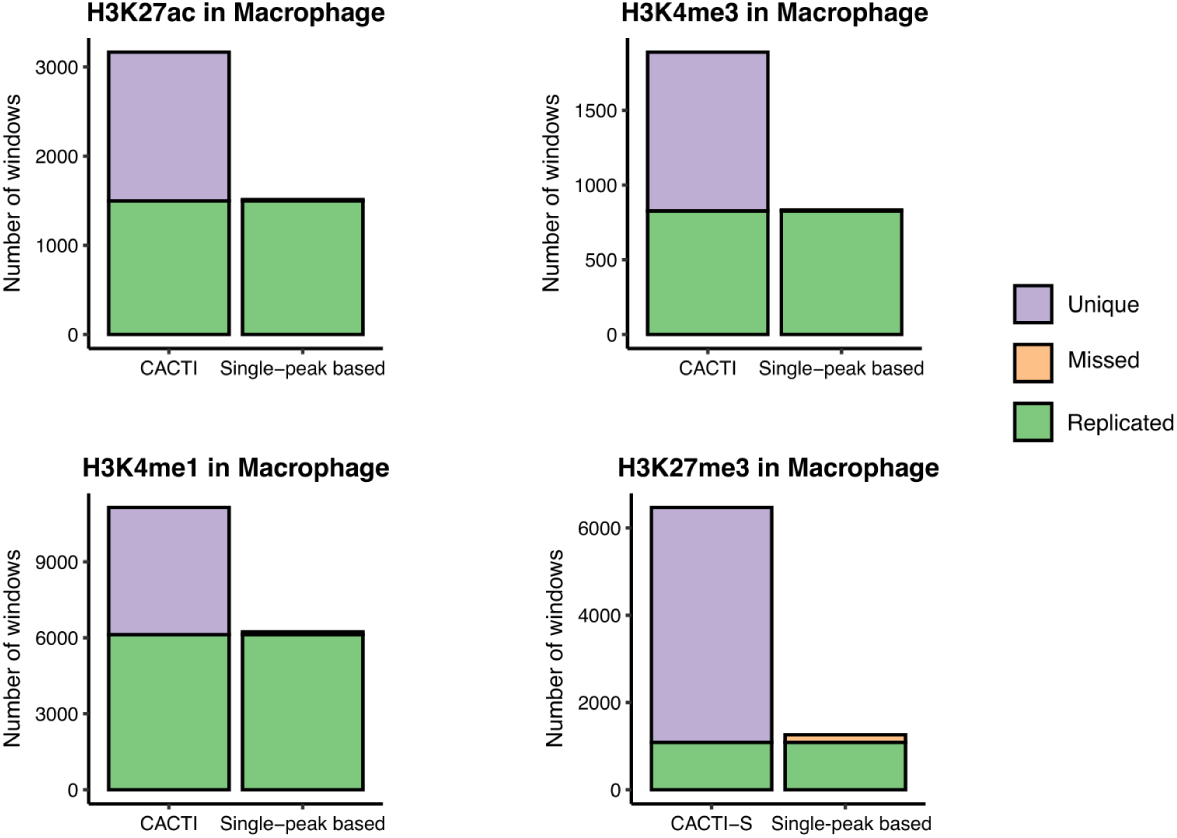
Performance comparison of CACTI to standard mapping methods for four marks in macrophage. Replicated cQTLs (i.e., cWindows containing at least one cPeak) are represented in green, cQTLs specific to standard mapping methods (i.e., missed by CACTI) are represented in yellow, and cQTLs Unique to CACTI (i.e., missed by standard methods) are represented in purple.

**Figure S11.**
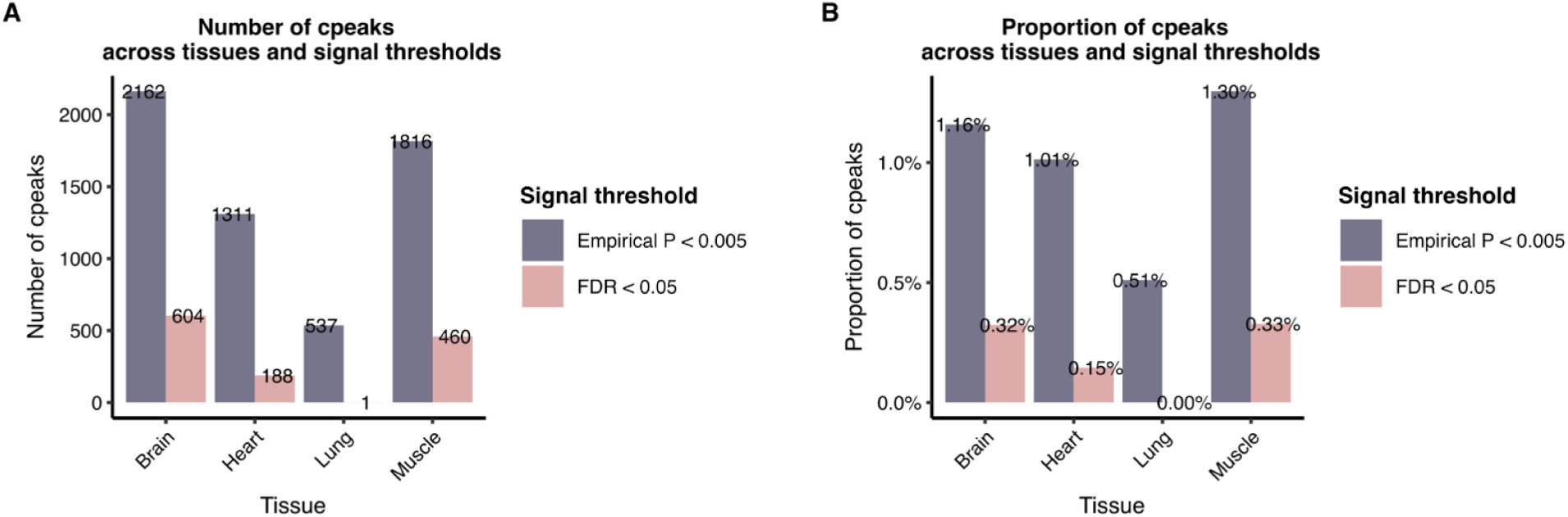
Number of cPeaks under different p-value cutoffs in eGTEx. X-axis gives the four tissues in eGTEx. Pink bars represent cPeaks using the FDR threshold lower than 0.05. We performed multiple testing correction on all peaks using q-value. Grey bars represent cPeaks using the p-value threshold used in the eGTEx paper, which is to use an empirical p-value threshold of 0.005. (A) Y-axis gives the number of cPeaks. (B) Y-axis gives the proportion of cPeaks among all peaks defined in each dataset.

**Figure S12.**
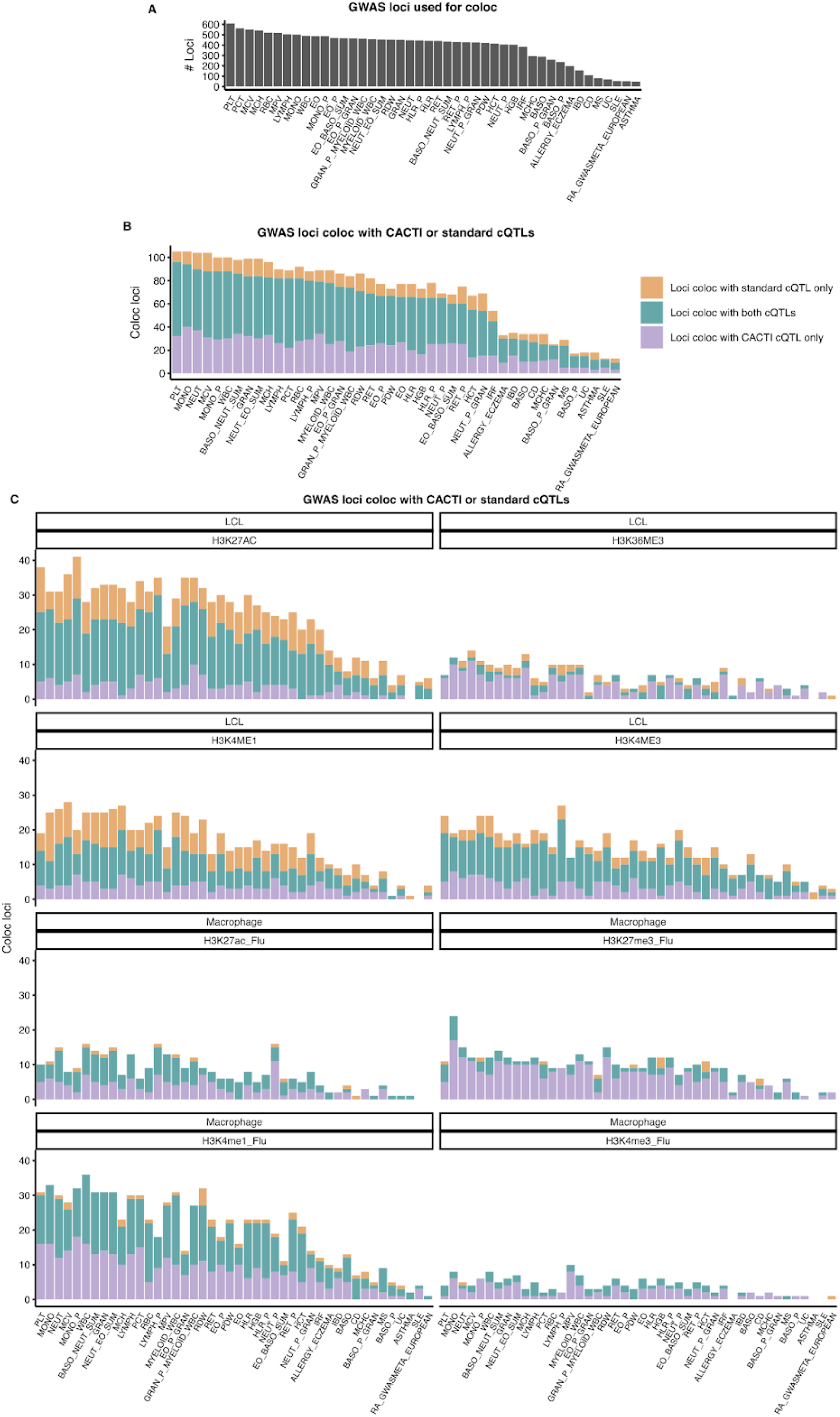
Colocalization of GWAS loci of 36 blood related traits and 8 immune diseases with cQTLs across marks and cell types. (A) Number of GWAS loci used to perform colocalization analysis for 44 traits. (B) GWAS loci coloc with cQTLs by standard mapping method or CACTI for any mark in any cell types. Purple bars show the loci colocalized with only CACTI cQTLs. Yellow bars show the loci that colocalize with only standard cQTLs. Blue bars show the loci colocalized with both types of cQTLs. (C) GWAS loci coloc with cQTLs in individual datasets. Each panel is a dataset.

**Figure S13.**
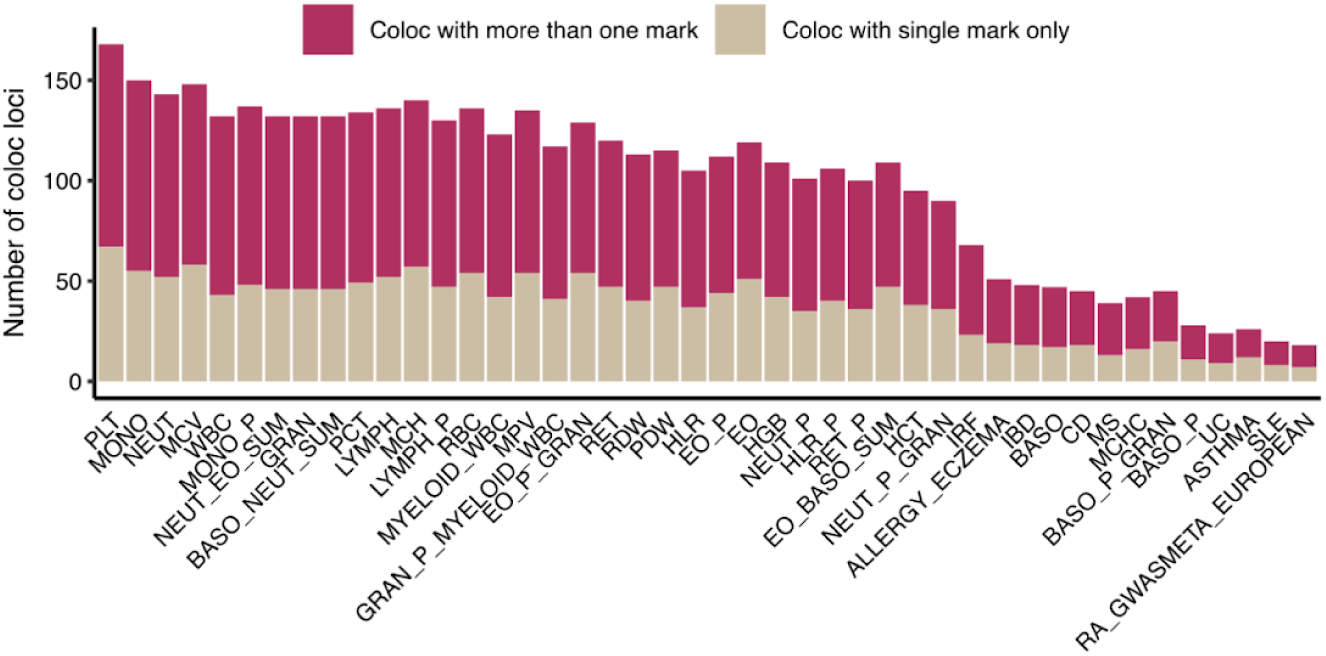
Sharing and specificity of GWAS loci coloc with cQTLs of individual marks. X-axis shows the 44 traits and y-axis shows the number of coloc loci. Yellow bars show the number of GWAS loci coloc with only one mark. Red bars show loci coloc with at least two marks. Marks H3K27ac, H4K4me1, H3K4me3, H3K36me3, and H3K27me3 were used.

**Figure S14.**
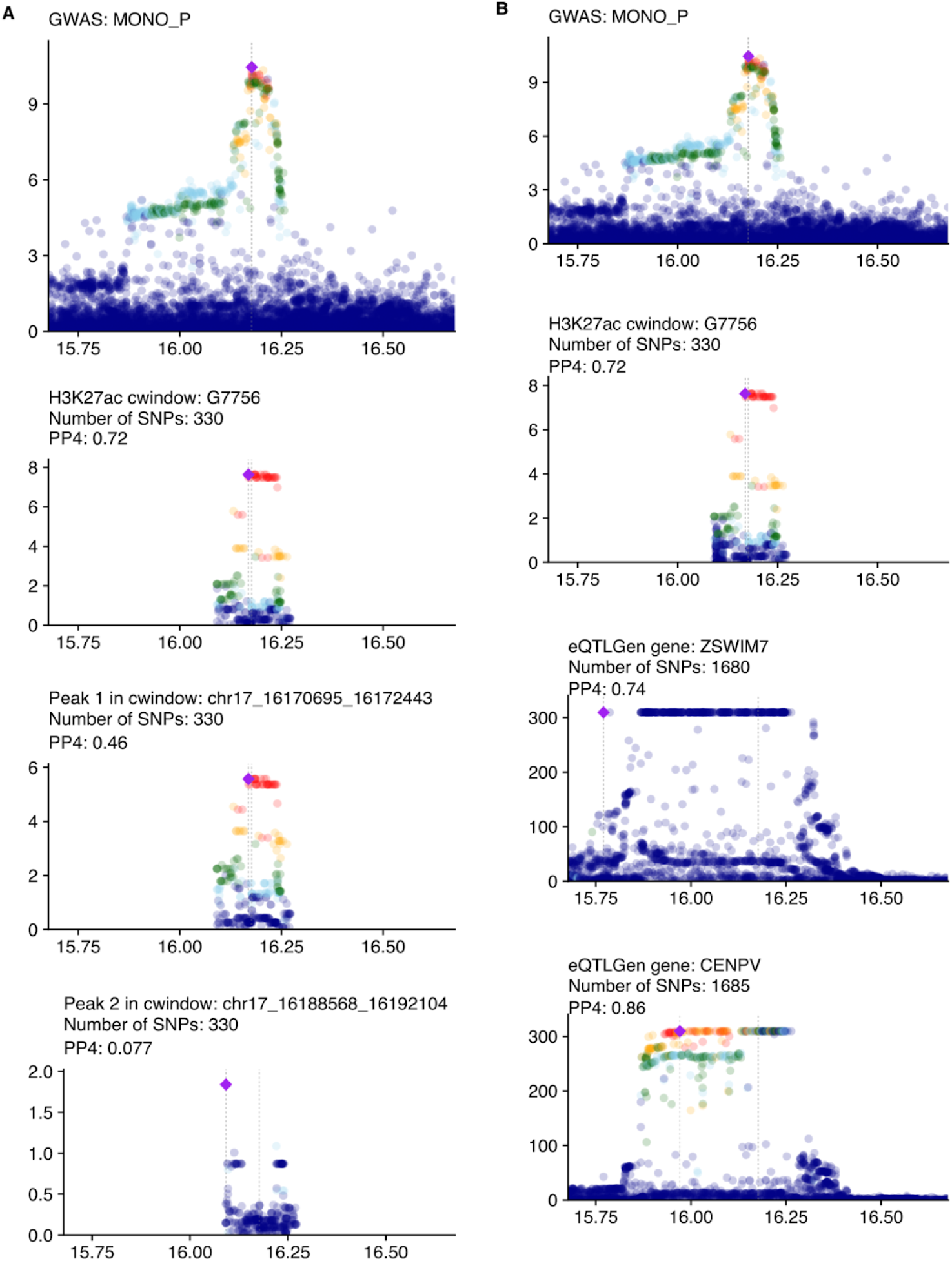
Manhattan plot for coloc example in. **Figure 3D**. The first row shows the GWAS locus of the example trait. The second row shows the CACTI association between the GWAS locus and cWindow in the example. The number of SNPs in the region and PP4 are shown in the title. (A) The remaining two rows give the univariate associations between the GWAS locus and cQTLs of two peaks contained in the cWindow. (B) The remaining two rows give the univariate associations between the GWAS locus and eQTLs of two genes nearby (*CENPV* and *ZSWIM7*).

**Figure S15.**
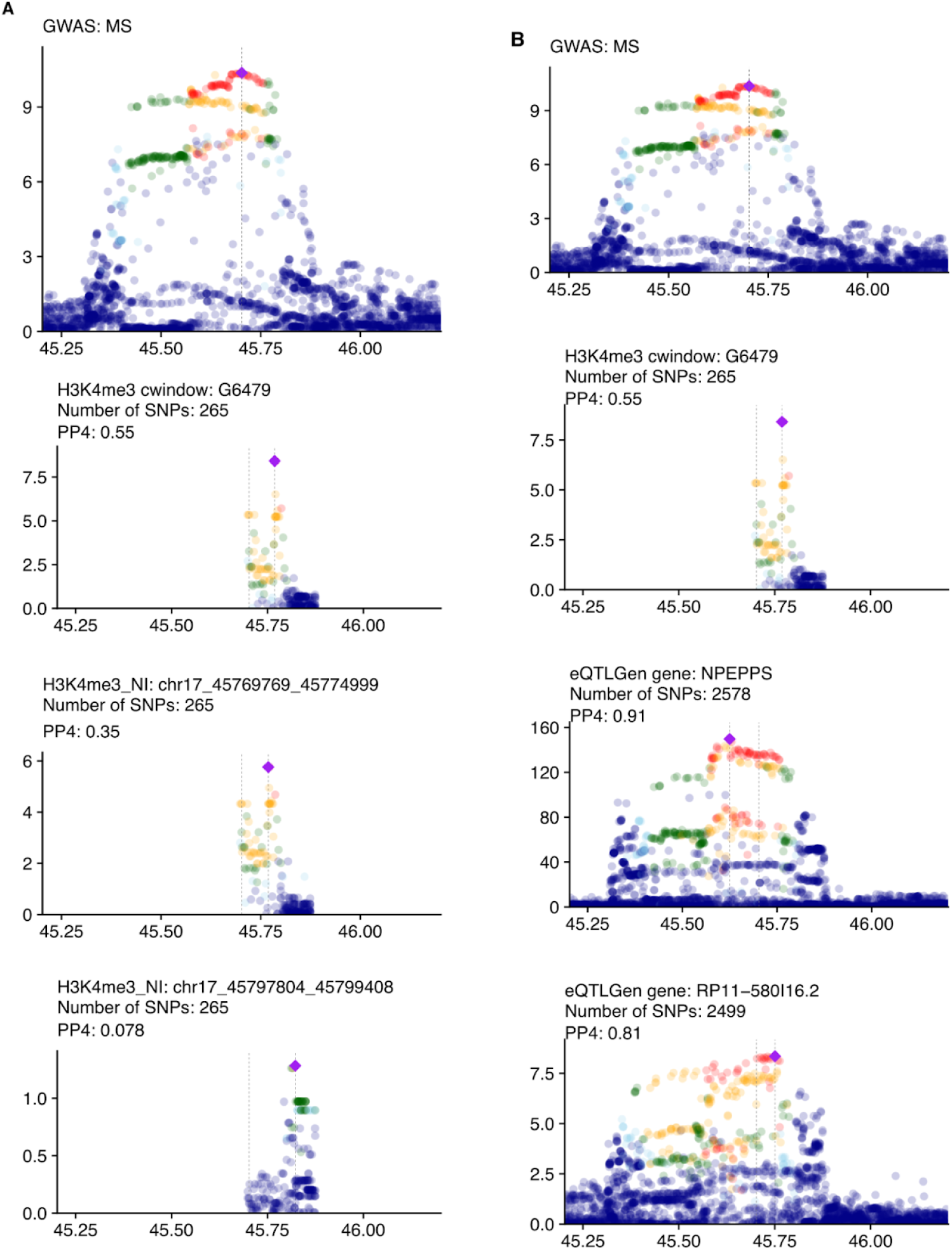
Manhattan plot for coloc example in. **Figure 3E**. The first row shows the GWAS locus of the example trait. The second row shows the CACTI association between the GWAS locus and cWindow in the example. The number of SNPs in the region and PP4 are shown in the title. (A) The remaining two rows give the univariate associations between the GWAS locus and cQTLs of two peaks contained in the cWindow. (B) The remaining two rows give the univariate associations between the GWAS locus and eQTLs of two genes nearby (*NPEPPS* and *RP11-580I16.2*).

**Figure S16.**
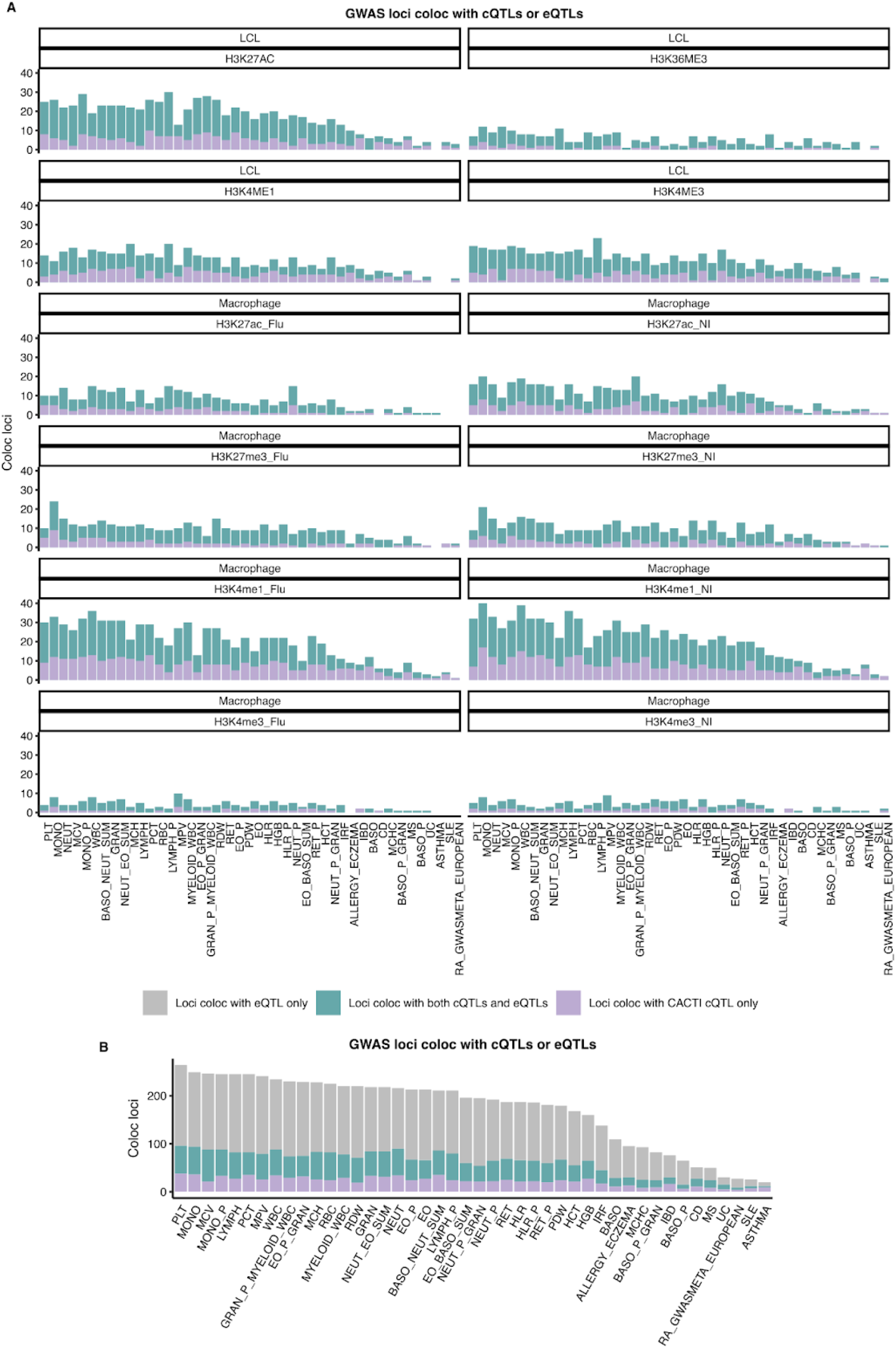
Colocalization of GWAS loci with CACTI cQTLs or eQTLGen eQTL. (A) Number of GWAS loci colocalized with CACTI cQTLs only (purple) or with both CACTI cQTLs and eQTLs (blue) across individual cQTL datasets. Each facet represents a dataset. Traits are shown on the x-axis. (B) Summary of GWAS loci colocalization using cQTLs from all datasets combined and all eQTLS from eQTLGen. Bars represent the number of loci colocalizing with CACTI cQTLs only (purple), with both CACTI cQTLs and eQTLs (blue), or with eQTLs only (grey), across traits on the x-axis.

**Figure S17.**
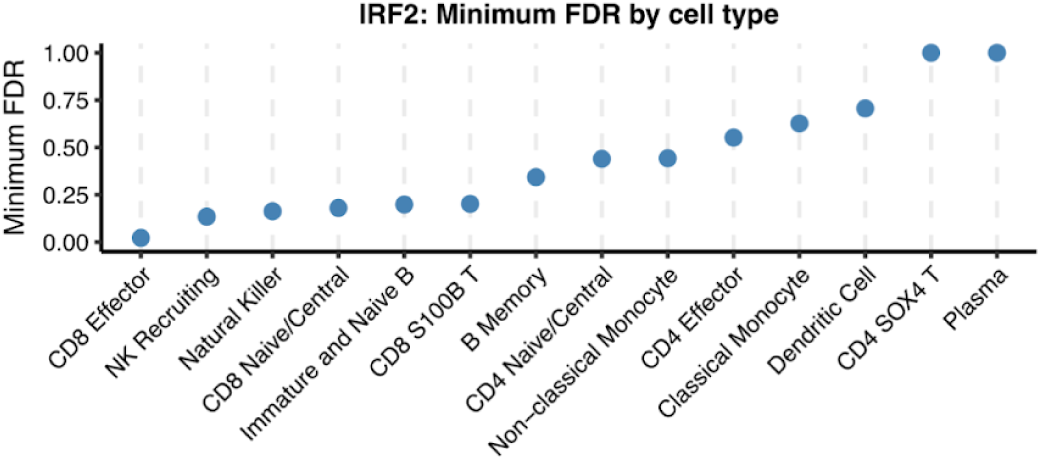
Minimum FDR across immune cell subpopulations in the OneK1K cohort. The scatter plot displays the minimum FDR of cell-type–specific eQTL for *IRF2* across 14 immune cell types in the OneK1K dataset.

### SUPPLEMENTAL TABLES

Table S1. Associations of all cWindows and lead SNPs across datasets.

Table S2. Enrichment of CACTI-specific cQTLs, all CACTI-cQTLs, and shared cQTLs between CACTI and single-peak-mappings for eQTLs.

Table S3. Enrichment of CACTI-specific cQTLs and all CACTI-cQTLs for eQTLs, using peak-number matched windows as backgrounds.

Table S4. Macrophage-relevant cell types used to calculate ABC links.

Table S5. Covariates number used in each dataset.

